# A stable leading strand polymerase/clamp loader complex is required for normal and perturbed eukaryotic DNA replication

**DOI:** 10.1101/808840

**Authors:** Katy Stokes, Alicja Winczura, Boyuan Song, Giacomo De Piccoli, Daniel B. Grabarczyk

## Abstract

The eukaryotic replisome must faithfully replicate DNA and cope with replication fork blocks and stalling, while simultaneously promoting sister chromatid cohesion. Ctf18-RFC is an alternative PCNA loader that links all these processes together by an unknown mechanism. Here, we use integrative structural biology combined with yeast genetics and biochemistry to highlight the specific functions that Ctf18-RFC plays within the leading strand machinery *via* an interaction with the catalytic domain of DNA Pol ε. We show that a large and unusually flexible interface enables this interaction to occur constitutively throughout the cell cycle and regardless of whether forks are replicating or stalled. We reveal that, by being anchored to the leading strand polymerase, Ctf18-RFC can rapidly signal fork stalling to activate the S phase checkpoint. Moreover, we demonstrate that, independently of checkpoint signaling or chromosome cohesion, Ctf18-RFC functions in parallel to Chl1 and Mrc1 to protect replication forks and cell viability.

## Introduction

The faithful maintenance of genetic information is essential for cell viability and regulated proliferation; strikingly, defects in chromosome duplication and segregation are a powerful source of genomic instability and a hallmark of cell transformation (Hanahan and Weinberg, 2011). DNA duplication depends on a complex machine, the replisome, which contains the DNA helicase CMG (Cdc45-Mcm2-7-GINS) that unwinds the parental DNA (Gambus *et al*., 2006; Ilves *et al*., 2010; Pacek *et al*., 2006), and three DNA polymerases, namely DNA Pol ε, α, and δ, the first required for the synthesis of the majority the leading strand and the latters for initial DNA synthesis following origin firing and the synthesis of the lagging strand (Burgers and Kunkel, 2017; Zhou *et al*., 2019). Both Pol ε and δ are processive DNA polymerases that interact with the PCNA sliding clamp (Chilkova *et al*., 2007). In addition, the replisome includes several other proteins, which provide both physical and functional coordination and regulation of DNA unwinding and DNA synthesis, such as, Ctf4 (Samora *et al*., 2016; Simon *et al*., 2014; Villa *et al*., 2016) and Mrc1 (Yeeles *et al*., 2017; Hodgson, Calzada and Karim, 2007; Tourrière *et al*., 2005).

During S phase, however, eukaryotic cells must not only faithfully duplicate their DNA, but also promote the association of the resulting sister chromatids, which must be kept together until the metaphase to anaphase transition. Moreover, replication forks must cope with DNA damage and stalling and promote the recruitment and activation of protein kinases, in a pathway called the S phase checkpoint (Labib and De Piccoli, 2011). Therefore, chromosome maintenance requires that DNA synthesis, chromosome cohesion establishment and the ability to activate the S phase checkpoint must occur simultaneously at the replication fork. Strikingly deletions of a single component of the replisome often can cause considerable defects in DNA synthesis, in chromosome cohesion (Borges *et al*., 2013) and activation of the S phase checkpoint kinase Rad53 in response to fork stalling (Alcasabas *et al*., 2001; Can *et al*., 2019), thus highlighting how DNA synthesis is closely intertwined with other processes necessary for the maintenance of genome stability. The replicative factor Ctf18-Dcc1-Ctf8-Rfc2-5 complex (Ctf18-RFC) exemplifies these overlapping functions at forks.

Ctf18-RFC belongs to the RFC family of AAA+ ATPase clamp loaders, which are involved in the loading and unloading of clamps onto dsDNA (Bylund and Burgers, 2005; Kouprina *et al*., 1994). *In vitro* analyses have shown that Ctf18-RFC loads PCNA, albeit with lower efficiency than the essential PCNA-loader RFC1 (Bylund and Burgers, 2005). In addition, Ctf18-RFC shows an ATP-dependent PCNA unloading activity (Bylund and Burgers, 2005), although it is still not understood whether this unloading occurs *in vivo* (Lengronne *et al*., 2006). Importantly, Ctf18-RFC localizes at replication forks during S phase and it is required for chromosome cohesion (Lengronne *et al*., 2006; Hanna *et al*., 2001). Mutants of each subunit show defects in cohesion and chromosome maintenance, decreased levels of Smc3 acetylation, and synthetic lethality in combination with replisome mutants defective in chromosome cohesion such as *CTF4* or *CHL1* (Mayer *et al*., 2001; Borges *et al*., 2013; Mayer *et al*., 2004; Petronczki *et al*., 2004). Moreover, Ctf18-RFC is required, together with Mrc1, for the activation of the S phase checkpoint (Kubota *et al*., 2011; Crabbé *et al*., 2010). In *ctf18Δ* cells, checkpoint activation in response to fork stalling following HU treatment is delayed and depends on the DNA damage checkpoint mediators Rad9 and Rad24 (Crabbé *et al*., 2010).

Compared to other clamp loaders, Ctf18-RFC contains two additional subunits, Ctf8 and Dcc1, which bind to the C-terminus of Ctf18 to form the Ctf18-1-8 module (Bylund and Burgers, 2005). This module is separated from the RFC catalytic module by a large predicted unstructured linker (Fig 1A). The Ctf18-1-8 module comprises a triple-barrel domain (TBD) formed from all three proteins and a winged helix hook (WHH), composed of a series of winged-helix domains in the C-terminus of Dcc1 resembling a hook (Grabarczyk, Silkenat and Kisker, 2018; Wade *et al*., 2017). Mutations in Ctf18 that disrupt formation of this Ctf18-1-8 module result in similar but not identical phenotypes as entire deletions of Ctf18 (Okimoto *et al*., 2016; García-Rodríguez *et al*., 2015), suggesting that this module is critical for at least some of the specialized functions of Ctf18-RFC. It has been shown that the Ctf18-1-8 module binds the N-terminal catalytic domain of Pol2 (García-Rodríguez *et al*., 2015; Murakami *et al*., 2010), the largest subunit of the leading strand Pol ε, and that this interaction increases the clamp loader activity of Ctf18-RFC (Fujisawa *et al*., 2017). The Pol2 catalytic domain (Pol2_CAT_) is connected by an unstructured linker to the remaining parts of Pol ε, which form an integral part of the CMGE (CMG-Pol ε) core replisome complex (Fig 1A) (Goswami *et al*., 2018; Zhou *et al*., 2017). It was recently shown that the last winged helix domain, WH3, of Dcc1 assumes a dual function: it can transiently interact with either DNA or a minimal construct of Pol2 (Grabarczyk, Silkenat and Kisker, 2018). However, despite a triple amino acid mutation within the WH3 domain that completely abolishes both interactions *in vitro*, surprisingly, deletion of WH3 *in vivo* only causes a small loss of Ctf18-RFC from replicating forks, while a cohesion defect is only seen when both WH2 and WH3 are deleted (Wade *et al*., 2017). Thus, it remains unclear what the Ctf18-1-8 module is targeting Ctf18-RFC to and under which conditions this occurs.

**Fig 1.**
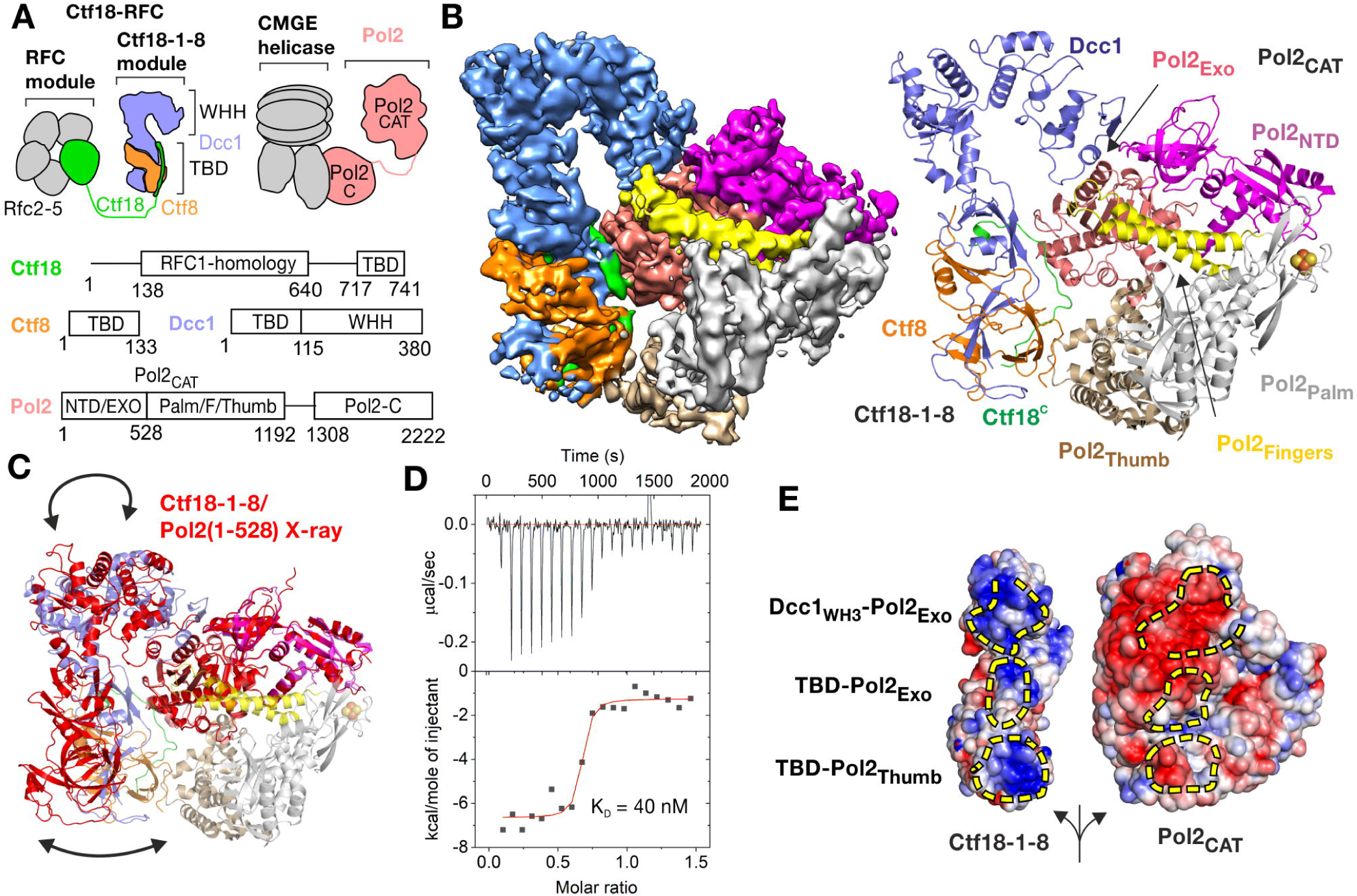
Cryo-EM structure of the Pol2_CAT_/Ctf18-1-8 complex. **(A)** Domain architecture and molecular context of Pol2 and Ctf18-RFC. CMGE refers to the Cdc45-Mcm2-7-GINS helicase in complex with the non-catalytic subunits of Pol ε. TBD – triple barrel domain, WHH – winged helix hook, NTD – N-terminal domain, F – fingers subdomain, C – C-terminal domain. **(B)** The left panel shows a sharpened cryo-EM map of the complex colored by Ctf18-1-8 subunit as in panel A and Pol2_CAT_ subdomain, with Pol2_NTD_ in magenta, Pol2_PALM_ in gray, Pol2_EXO_ in salmon, Pol2_FINGERS_ in yellow and Pol2_THUMB_ in brown. The right panel shows the refined model of the complex in cartoon representation with the iron-sulfur cluster as spheres. **(C)** Superposition by Pol2_NTD_-Pol2_EXO_ of the cryo-EM structure (colored as in panel B) and the Pol2(1-528)/Ctf18-1-8 crystal structure (pdb 5oki) shown entirely in red. **(D)** ITC experiment showing titration of 170 µM Ctf18-1-8 into 25 µM Pol2CAT in the presence of 50 µM P1P2. **(E)** The three interaction patches are shown on the opened surface of the complex as yellow dashed areas. The electrostatic surface was calculated using APBS and shown as a gradient from −4 kT/e (red) to +4 kT/e (blue).

To resolve these ambiguities, we have examined the interaction between Ctf18-1-8 and the entire catalytic domain of Pol2 using a combination of structural biology and yeast genetics. We show that the interaction has a nanomolar affinity, does not interfere with the polymerase functions of Pol2_CAT_, and occurs constitutively throughout the cell cycle, which together strongly suggests that Ctf18-RFC forms a stable part of the replisome leading strand machinery. Furthermore, we have solved the cryo-EM structure of this complex, which reveals a remarkably flexible interface that permits high-affinity binding of Ctf18-1-8 to different conformational states of Pol2_CAT_. Importantly, our structure has enabled us to identify residues that, when mutated, specifically abrogate the interaction between Ctf18-1-8 and Pol2 *in vivo*, both in yeast and human cells, allowing us to dissect the role of the Ctf18-RFC/Pol ε interaction. We show that an intact leading strand polymerase/clamp loader complex is essential for the replication stress checkpoint, but not cohesion establishment. This separation of function permits us to resolve the cohesion-independent roles of Ctf18-RFC, Mrc1 and Chl1 in DNA replication.

## Results

### Architecture of the Pol2_CAT_/Ctf18-1-8 complex

Previously, we showed that yeast Ctf18-1-8 forms a relatively transient complex with a truncated construct of the Pol ε catalytic domain, Pol2(1-528) (Fig, 1A), through the WH3 domain of Dcc1 (Dcc1_WH3_) and exonuclease domain of Pol2 (Pol2_EXO_) (Grabarczyk, Silkenat and Kisker, 2018). Although this minimal complex could be disrupted *in vitro* by a structure-guided triple mutation, Dcc1 R367A/R376A/R380A (henceforth Dcc1-3A) (Grabarczyk, Silkenat and Kisker, 2018), a new crystal form of this complex revealed that this interface is highly flexible (Figure S1A). We therefore used cryo-EM to solve the structure of Ctf18-1-8 in complex with the complete catalytic domain of Pol2 (residues 1-1192, henceforth Pol2_CAT_) to 4.2 Å resolution. This is the maximal part of the Pol ε/Ctf18-RFC complex that can be structurally analyzed due to the large-scale flexibility of both Pol ε and Ctf18-RFC (Fig. 1A) (Shiomi *et al*., 2004; Zhou *et al*., 2017).

Our structure reveals that the complete Ctf18-1-8/Pol2_CAT_ complex has a large interface (Fig. 1B) burying an area of 1363 Å^2^, which is more than double the 612 Å^2^ area buried in the minimal complex. Dcc1_WH3_ interacts with Pol2_EXO_ as previously observed, but this interface is now further stabilized by the Pol2 fingers subdomain (Pol2_FINGERS_). Furthermore, entirely new interactions are formed by the Ctf18-1-8 triple-barrel domain (TBD), which has rotated towards the polymerase (Fig. 1C) to interact with Pol2_EXO_ and the Pol2 thumb (Pol2_THUMB_) subdomain in two discontinuous interaction patches (Fig. S1B). The large interface suggested that this might be a stable interaction, and so we measured the affinity between Ctf18-1-8 and Pol2_CAT_ by isothermal titration calorimetry. This showed that Ctf18-1-8 forms a tight complex with Pol2_CAT_ with a *K*_*D*_ of 40 nM (Fig. 1D), over 30-fold stronger than the 1.3 µM *K*_*D*_ measured for the truncated Pol2(1-528) construct (Grabarczyk, Silkenat and Kisker, 2018).

Unlike the Dcc1_WH3_-Pol2_EXO_ and TBD-Pol2_THUMB_ interfaces, which are mostly electrostatic in nature (Fig. 1E) and show few specific interactions, the TBD-Pol2_EXO_ patch is well-ordered and displays hydrophobic and backbone interactions involving highly-conserved amino acids (Fig. 2A). This interface centers on an extension of the Pol2_EXO_ beta-sheet into the upper beta-barrel of Ctf18-1-8, *via* a beta-strand formed by the absolutely conserved ^730^V(R/K)(R/K)^732^ motif in Ctf18, resulting in a distorted four-protein anti-parallel beta-sheet (Fig. 2A). This interaction is further stabilized by the burying of Ctf18 Val730 into a hydrophobic pocket in Pol2_EXO_ formed by Phe335, Tyr337 and Val475 (Fig. 2A). To test the importance of this interface for stabilizing the complex, we generated a Ctf18 V730R/R731A/K732A variant (henceforth Ctf18-RAA). In an EMSA supershift assay, wild-type Ctf18-1-8 saturated Pol2_CAT_-DNA at the lowest tested concentration of 125 nM (Fig. 2B), while the Dcc1-3A and Ctf18-RAA variants showed a clear binding defect. A variant combining both was completely disrupted for Pol2_CAT_ binding under the tested conditions (Fig. 2B), validating the structure.

**Fig 2.**
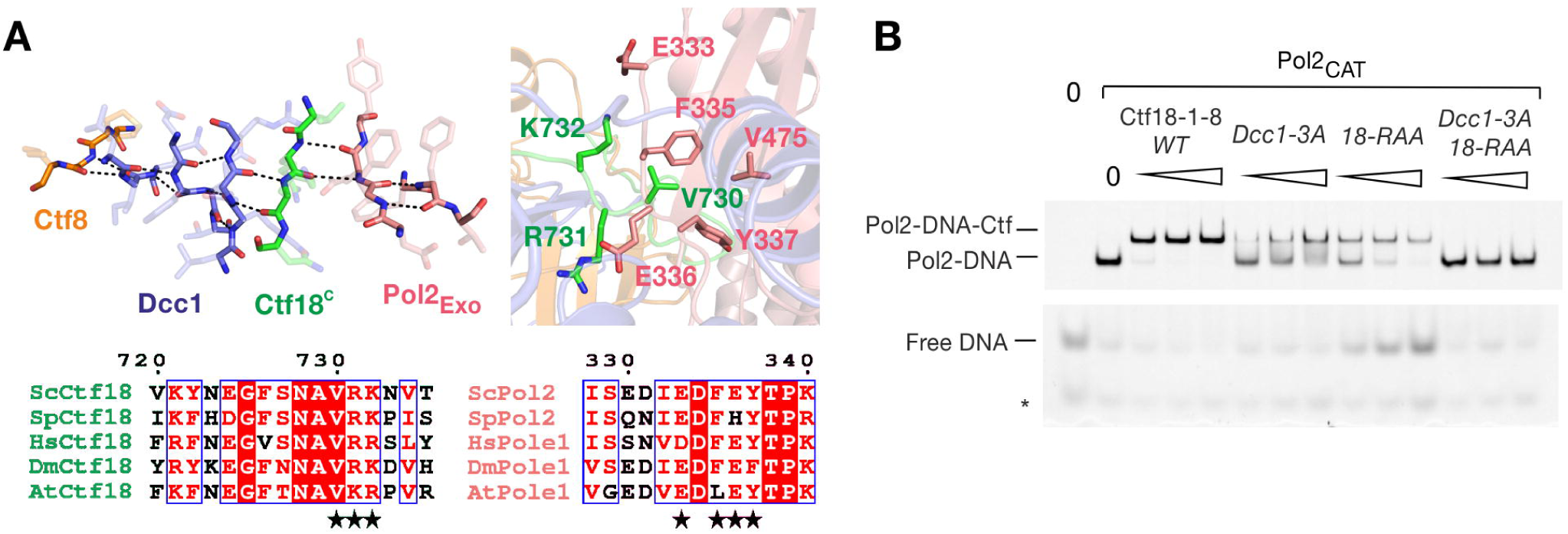
The highly-conserved TBD-Pol2_EXO_ interface stabilizes the Ctf18-1-8/Pol2_CAT_ complex. **(A)** Molecular details of the TBD-Pol2_EXO_ patch, showing backbone interactions in the left panel, and selected sidechain interactions in the right panel, with coloring as in Fig. 1B. Alignments were performed in ClustalX2 and displayed using ESPript 3.0. Sc – *Saccharomyces cerevisiae*, Sp – *Schizosaccharomyces pombe*, Hs – *Homo sapiens*, Dm – *Drosophila melanogaster*, At – *Arabidopsis thaliana*. **(B)** EMSA supershift assay using 50 nM P1P2, 125 nM Pol2_CAT_ and 125, 250 and 500 nM of the indicated Ctf18-1-8 variants.

### Plasticity of the Pol2_CAT_/Ctf18-1-8 interaction

Comparing our DNA-free structure of Pol2_CAT_ with the crystal structure of the DNA- and nucleotide-trapped Pol2_CAT_ shows movements of the Pol2_THUMB_, Pol2_PALM_ and Pol2_FINGERS_ subdomains that are characteristic of B-family DNA polymerases (Fig. S2C) (Franklin, Wang and Steitz, 2001). As Ctf18-1-8 binds across multiple subdomains that should move during DNA binding and catalysis, it might be expected that the interaction inhibits the polymerase. Remarkably, when we tested the activity of Pol2_CAT_ in a primer extension assay, the presence of Ctf18-1-8 had no effect on activity (Fig. 3A). Furthermore, analysis of DNA substrate engagement using an EMSA, showed identical DNA binding properties for Pol2_CAT_ and the Pol2_CAT_/Ctf18-1-8 complex (Fig. 3B). This suggests that, despite the large interface, the interaction has no direct role in regulating the behaviour of the polymerase. We next asked the question from the inverse perspective – would DNA engagement by the polymerase affect the interaction between Ctf18-1-8 and Pol2_CAT_? Strikingly, ITC showed a nearly identical *K*_*D*_ for the interaction in either the presence or absence of the polymerase substrate (Fig. S1D), suggesting that the complex is stable independent of Pol2 function.

**Fig 3.**
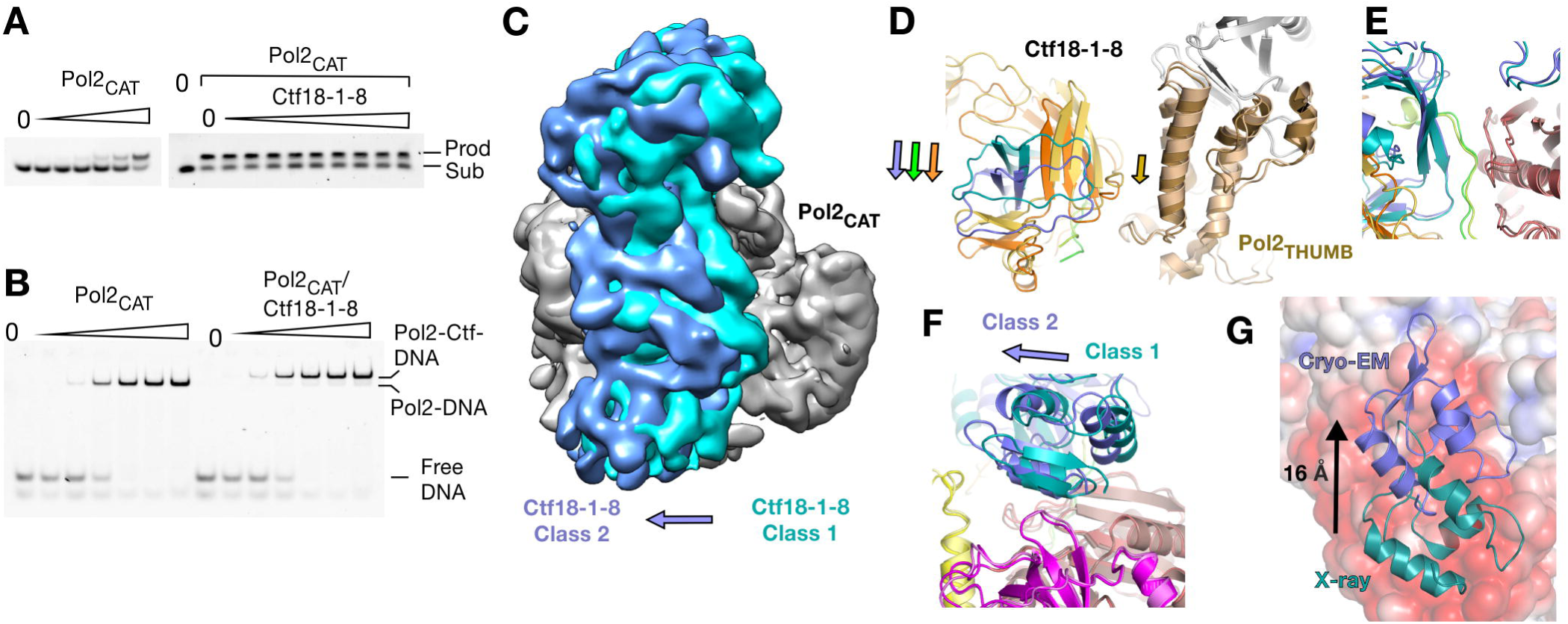
Plasticity of the Ctf18-1-8/Pol2_CAT_ interaction. **(A)** Primer extension assay of an 11 base primer with a 16 base template (P1P2) with increasing Pol2_CAT_ in a two-fold dilution series from 0.09 nM to 3 nM in the left panel, and in the right panel a constant Pol2_CAT_ concentration of 1 nM with a two-fold dilution series of Ctf18-1-8 from 8 nM to 2 µM. **(B)** Interaction EMSAs using the P1P2 substrate, with a two-fold dilution series from 0.125 to 2 µM of either Pol2_CAT_ alone or a 1:1 Pol2_CAT_/Ctf18-1-8 complex. **(C)** A superposition of Class 1 (teal) and Class 2 (blue) by their Pol2_NTD-EXO_ domains. Class 1 is Gaussian filtered to a similar resolution as Class 2 for clarity. **(D-F)** Superposition of the models from cryo-EM classes 1 and 2, aligned by Pol2_NTD-EXO._ **(D)** Corresponding movements of Pol2_THUMB_ and the TBD (**E**) The TBD-Pol2_EXO_ interface is unchanged between Class 1 and Class 2. **(F)** Dcc1_WH3_ forms different interactions with Pol2_EXO_ in each class. **(G)** Superposition of the Pol2(1-528)/Ctf18-1-8 complex and Pol2_CAT_/Ctf18-1-8 complex as in Fig. 1C but the Pol2_CAT_ surface is colored by charge with the same parameters as Fig. 1E and Dcc1 is colored in teal for the X-ray structure and blue for the Cryo-EM structure (Class 1).

We next investigated the mechanism by which the large Ctf18-1-8 binding interface does not impose thermodynamic penalties onto the conformational changes of Pol2_CAT_. Although the *K*_*D*_ values were identical, our ITC showed a clear difference in *ΔH* in the presence and absence of the DNA substrate (Fig. S1D), suggesting that the interaction mechanism is distinct in each case. As glutaraldehyde fixation was necessary to obtain sufficient sample quality for structure determination, it was not straightforward to obtain a catalytically engaged structure of the complex. Instead, we could obtain clear insight into how the interaction behaves in different Pol2_CAT_ conformations by refining a second lower-resolution (5.8 Å) cryo-EM class (Class 2) (Fig. 3C). In Class 2, while the majority of Pol2_CAT_ remains in the same position, Pol2_THUMB_ has undergone a small movement (Fig. 3D). To maintain simultaneous interaction at all three patches despite this movement, the entirety of Ctf18-1-8 has rotated (Fig. 3C-F). While the central highly specific interface is unchanged (Fig. 3E), the Dcc1_WH3_-Pol2_EXO_ interaction is completely altered (Fig. 3F). This plasticity can be explained by the non-specific and electrostatic nature of this interface. The crystal structures provide an extreme example of this. Here, Dcc1_WH3_ has slid by 16 Å along the negatively charged Pol2_EXO_ surface (Fig. 3G) and forms entirely different molecular interactions with it (Fig. S1E). Although this conformation is presumably influenced by crystal contacts, approximately the same interface is utilized in two different crystal forms, suggesting it is a readily accessible state in the conformational continuum. Thus, our data show that the complex uses slippery electrostatic interfaces with low specificity that are readily adaptable to maintain tight complex formation in different conformational states.

### Ctf18-RFC is recruited to replication forks by Pol2

Our biochemical data have resolved the molecular details of the interaction between Pol2_CAT_ and Ctf18-1-8, and indicate that they form a constitutive complex. Next, we tested if this was true in the context of the replisome and replication forks in cells. First, we wanted to understand whether the interaction between Ctf18-RFC and Pol ε occurred *in vivo* throughout the cell cycle or only at replication forks. To this aim, synchronized cell cultures were treated with formaldehyde at different points of the cell cycle and Ctf18 was immunoprecipitated under stringent conditions. We observed that the interaction was crosslinking-dependent (Fig S4A), thus excluding the possibility of *ex vivo* binding. Interestingly, Ctf18-RFC and Pol ε interact constitutively throughout the cell cycle, independently of the presence of replication forks (Fig 4A). Next, we analyzed whether the binding between Ctf18-RFC and Pol ε depends on the interface described above. We first performed co-immunoprecipitation experiments with yeast strains containing a series of structure-guided mutations introduced at the genomic locus predicted to disrupt the interaction between Ctf18-RFC and Pol ε (Fig. 4B,C; S4B,C). We observed that most of the mutations, with the exception of *dcc1-3A*, negatively affected complex formation. To further disrupt the interaction between Ctf18-RFC and Pol ε, we combined *ctf18-RAA* either with *pol2-5A* (mutated for the acidic patch in Pol2_EXO_) or *dcc1WH3Δ*. The double mutant strains were mildly sensitive to drugs which induce fork stalling, *i*.*e*. the ribonucleotide reductase inhibitor hydroxyurea (HU), the DNA alkylating agent methyl methanesulphonate (MMS) and dsDNA-break inducer bleomycin, albeit to a lower extent than *ctf18Δ* (Fig. 4D). To ensure that the mutations did not affect the stability of the Ctf18-RFC complex itself, we conducted a mass spectrometry analysis and observed no difference in their composition (Fig. 4E). To test whether the loss of interaction between Ctf18-RFC and Pol ε leads to the displacement of these complexes from forks, we analysed their association to Mcm3 during DNA replication following cross-linking, thus maintaining transient protein-protein or protein-DNA interactions. Strikingly, while Pol2 and PCNA binding to Mcm3 were not affected in the mutant backgrounds, Ctf18-RFC was largely lost from the replisome (Fig. 4F). The interaction between Mcm3 and Ctf18-RFC depended on origin firing, since depletion of the replication initiation kinase DDK blocked Ctf18-RFC binding to Mcm3 (Fig. S4D), and it is maintained following replication stress (Fig S4E). Finally, we tested whether the complex has the same architecture in human cells. We observed that, while the transient expression of N- or C-terminal GFP-tagged alleles of POLE1 was able to co-immuno-precipitate Ctf18, the *pole1-5A* mutation (analogous to *pol2-5A*) abolished the interaction between the two complexes independently of the tagging orientation, suggesting that the acidic patch in POLE1_EXO_ is important for the recruitment of CTF18-RFC throughout evolution (Fig. 4G, S4F).

**Fig 4.**
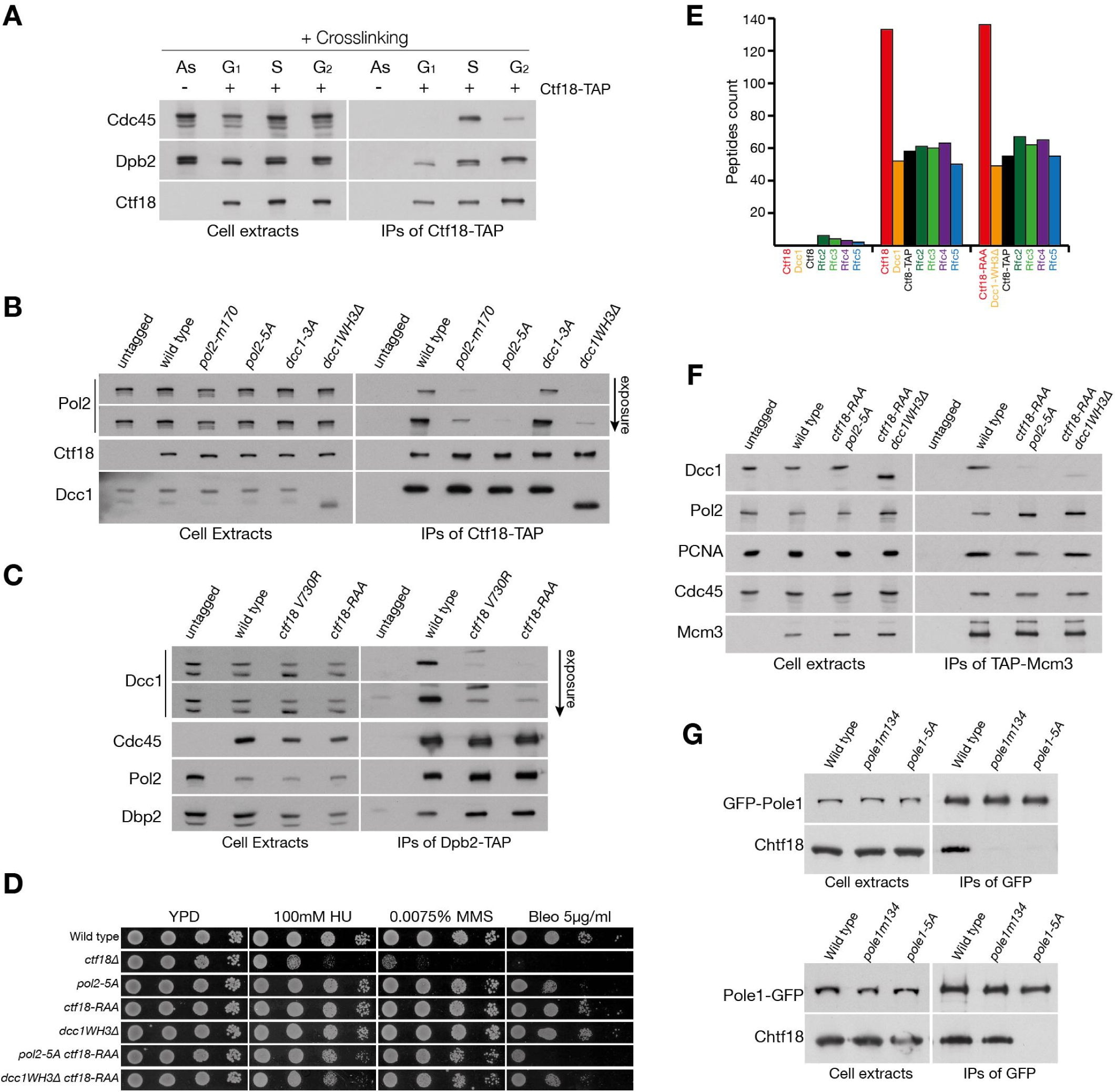
Ctf18-RFC recruitment at forks depend on its binding to Pol2. **(A)** Analysis of the interaction of Ctf18-RFC and Pol ε *in vivo*. Cells, carrying a TAP-tagged or untagged version of *CTF18* were grown to exponential phase (As) and arrested in G1, before being released into the cell cycle for 30 (S) or 60 minutes (G2). All cultures were treated with formaldehyde. The cross-linked protein extracts were then incubated with anti-TAP beads. Cell extracts and IPs were then analyzed by immunoblotting **(B**,**C)** Analysis of mutations in *POL2, CTF18* and *DCC1* on the Pol ε/Ctf18-RFC interaction. All mutants were generated at the genomic locus. These are *pol2* 170-GRAAAATGDAAG-181 (*pol2 m170*), *pol2 330-AAIAAFA-336* (*pol2-5A*), *dcc1 K364A, K367A, K380A* (*dcc1-3A*) and *dcc1(1-318)* (*dcc1WH3Δ*). *pol2-m170* was selected for analysis following a screen by Yeast-Two-Hybrids of conserved amino acids predicted to be on the surface of the protein (Fig. S4B-C). **(B)** Strains carrying a TAP-tagged allele of Ctf18, or an untagged control, were grown to exponential phase. Cell extracts and proteins immunoprecipitated using anti-TAP beads were analyzed by immunoblotting. **(C)** Wild type, *ctf18 V730R, or ctf18* 730-RAA-732 (*ctf18-RAA)* strains, carrying a *DPB2* or a *DPB2-TAP* allele, were analyzed as above. **(D)** Combination of double mutations causes replication stress sensitivity. The indicated strains were diluted 1:10 and spotted on the specified medium. **(E)** Mass spectrometry analysis of the Ctf18-RFC complex in wild type and *ctf18-RAA ddc1WH3Δ*. Cells carrying a TAP-tagged or untagged allele of *CTF8* were grown to the exponential phase before collection. Cell extracts were incubated with TAP-beads and washed at high salt (300 mM potassium acetate) before being analysed by mass spectrometry. The peptides reads for the CTF18-RCF complex are shown. **(F)** Loss of interaction with Pol2 causes the displacement of Ctf18 from replication forks. Cells carrying a TAP-tagged allele of *MCM3* and an untagged control were synchronously released in S phase and treated with formaldehyde. Cells extracts and the proteins immunoprecipitated with anti-TAP beads were analyzed by immunoblotting. **(G)** The interaction surface for the interaction between Pol ε and Ctf18-1-8 is conserved in human cells. HeLa cells were transiently transformed with plasmid expressing a N-terminal (top) or C-terminal (bottom) GFP-tagged alleles of *POLE1*, either wild type, *pole1 134-GPAAAADGPAAG-145* (*pole1-m134*), or *pole1 315-AAIAAFA-32*1 (*pole1-5A*). Cells extracts were incubated with anti-GFP beads, immunoprecipitated and analyzed by immunoblotting.

### Breaking the Pol ε/Ctf18-RFC interaction does not affect chromosome cohesion

With a molecular-level understanding of the physiological interaction, we were now able to determine which functions of Ctf18-RFC require it to be tightly tethered to the leading strand through an interaction with Pol ε. Ctf18-RFC is required for the establishment of chromosome cohesion in eukaryotic cells (Hanna *et al*., 2001; Mayer *et al*., 2001; Terret *et al*., 2009). Cells, carrying an mCherry-tagged *SPC42* allele and tetO repeats inserted at the *URA3* locus (Michaelis, Ciosk and Nasmyth, 1997), were synchronously released in S phase and mitotic cells were analysed for the number of foci present in the cell. We observed that both double mutants (*ctf18-RAA pol2-5A* and *ctf18-RAA dcc1WH3Δ*) showed no defects in cohesion (Figure 5A,B), while *ctf18Δ* showed defects in cohesion similar to that previously described (Xu, Boone and Brown, 2007). We also saw no sensitivity of the interaction mutants to benomyl, a drug that induces depolymerisation of microtubules, both in the presence or absence of the spindle checkpoint component *MAD2*. (Fig. 5C).

**Fig 5.**
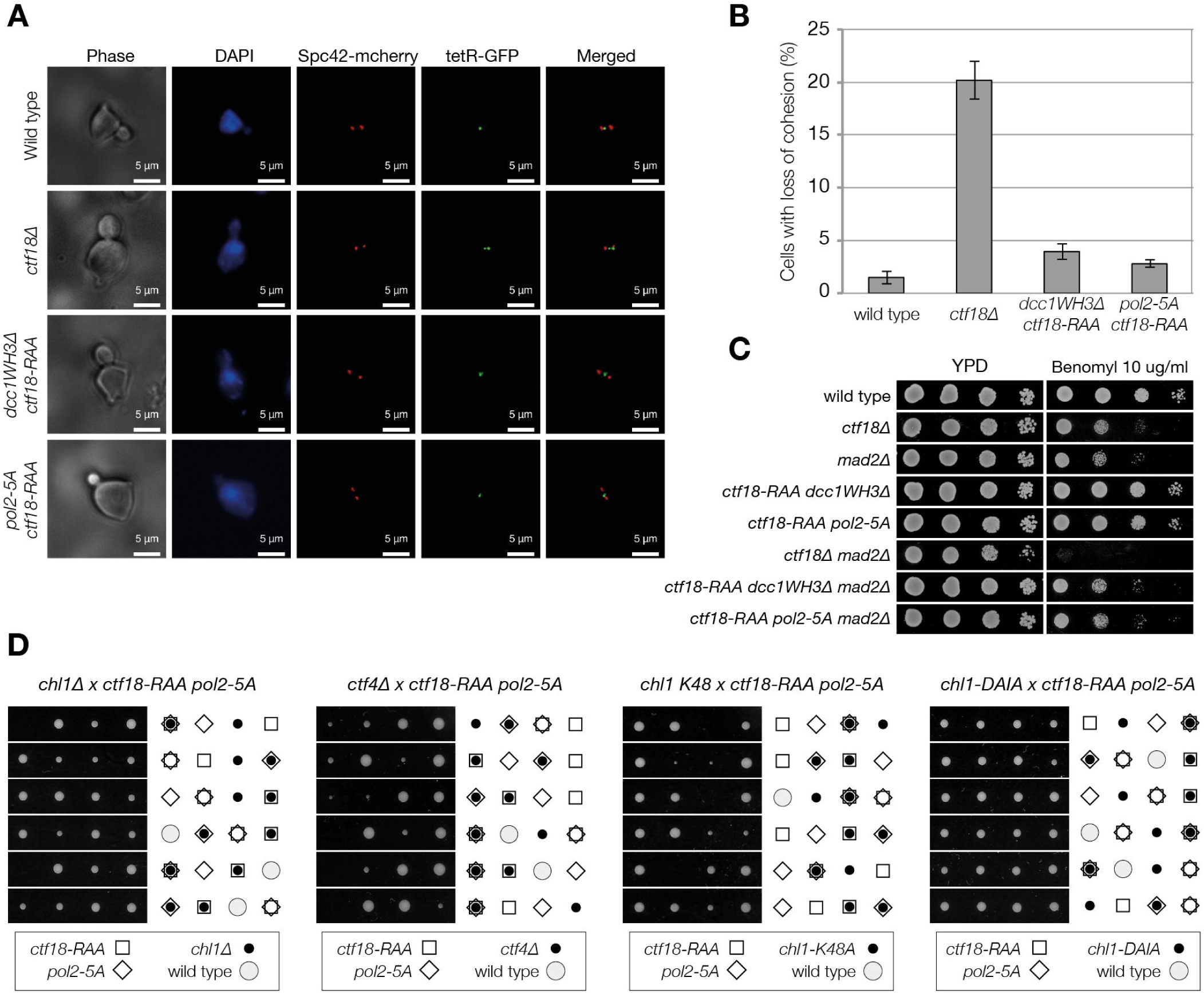
Breaking the interaction between Pol ε and Ctf18-RFC does not affect chromosome cohesion. **(A-B)** Analysis of chromosome cohesion. Cells carrying a *TetO* at the *URA3* locus (Michaelis, Ciosk and Nasmyth, 1997), a *TetR-GFP* and a *SPC42-mCHERRY* allele were arrested in G1 and synchronously released in S phase and collected ever 15 minutes. The distance between the Spc42-mCHERRY was scored and cells with spindles 1-2.5 µm were analyzed for number of GFP foci present. **(A)** Example of the images analyzed. **(B)** Summary of the results of three experiments. For each experiment and strain, around 400 cells with spindles 1-2.5 µm were analyzed. The average and standard deviation of three independent experiments are shown. **(C)** *ctf18-RAA pol2-5A/dcc1WH3Δ* are not sensitive to Benomyl nor show synthetic defects with *mad2Δ*. The strains were diluted 1:10 and spotted on the indicated medium. **(D)** Analysis of the meiotic progeny of diploids heterozygotes of the indicated alleles. Plates were scanned after 3 days growth at 24°C.

Next, we explored whether the interaction between Pol ε and the Ctf18-RFC complex is essential in the absence of other replication proteins required for cohesion establishment. *CTF4* and *CHL1* deletions cause defects in chromosome cohesion maintenance that are synthetic defective with CTF18-RFC mutations and are lethal in a *ctf18Δ* background (Hanna *et al*., 2001; Petronczki *et al*., 2004; Skibbens, 2004). Despite the lack of cohesion defects in our strains, surprisingly, we observed that *ctf18-RAA pol2-5A* is synthetic lethal in the absence of *CTF4* and *CHL1*. Chl1 is a DNA helicase that is recruited to forks through an interaction with Ctf4, and, as well as being required for chromosome cohesion, also has an important role during replication stress (Samora *et al*., 2016). To distinguish whether the synthetic lethality observed in the double mutant is a consequence of defects in DNA replication or loss of cohesion, we tested the genetic interaction between *ctf18-RAA po2-5A* and the allele *chl1-DAIA*, (which is deficient in chromosome cohesion) and the cohesion proficient but replication stress sensitive catalytic-dead allele *chl1 K48A* (Samora *et al*., 2016). Strikingly, we observed synthetic lethality with *chl1 K48A* but not with *chl1-DAIA*. Moreover, we observed that the deletion of the PCNA-interaction motif (PIP) at the C-terminus of Chl1 caused synthetic lethality/poor growth in a *ctf18-RAA pol2-5A* background (Figure S5A,B). We therefore conclude that loss Ctf18-RFC at the leading strand does not affect chromosome cohesion but causes defects at forks that require the action of Chl1 for the maintenance of cell viability.

### Cell viability and DNA replication depends on Mrc1 in cells lacking Ctf18-RFC at forks

To further test the function of the interaction, we screened for genetic interaction with various genes involved in DNA replication and repair and synthetic defective in a *ctf18Δ* background. We observed that the loss of the Ctf18-RFC/Pol ε interaction caused mild growth defects and DNA damage sensitivity in combination with *RAD52, HPR1, SGS1, SRS2, POL32* and *MRE11* (Fig. S5C,D). Strikingly, we observed synthetic lethality when MRC1 was deleted in *ctf18-RAA dcc1WH3Δ/pol2-5A* mutant backgrounds. This was not linked to defects in cohesion, since instead deleting *RMI1*, a gene epistatic to *MRC1* for chromosome cohesion (Lai *et al*., 2012), resulted in no defects (Fig. 6C). Mrc1 plays an important role in replication fork progression and re-start following fork stalling (Yeeles *et al*., 2017; Tourrière *et al*., 2005; Hodgson, Calzada and Karim, 2007). To analyse whether the genetic interaction of Mrc1 and our interaction mutants is related to a DNA replication defect, we introduced an auxin-inducible-degron Mrc1 allele and followed a single cell cycle. We observed that, while *the mrc1-aid ctf18-RAA dcc1WH3Δ/pol2-5A* triple mutants duplicated their DNA with similar kinetics to *mrc1-aid* cells, they activated the DNA damage checkpoint following the bulk of chromosome duplication (75 min) and arrested in G2/M as large budded cells by the end of the experiment (150 min) (Fig. 6D,E). Our data thus indicate that, the absence of Mrc1 and Ctf18-RFC at forks during DNA replication leads to an accumulation of DNA damage following DNA synthesis. Importantly, *mrc1-aid ctf18-RAA dcc1WH3Δ/pol2-5A* triple mutants show no additive defects in the activation of Rad53 in response to fork stalling, suggesting that the genetic interaction does not relate to checkpoint defects (Fig. 6F). Instead, our data suggest that loss of Ctf18-RFC from the leading strand results in a DNA replication defect that is compensated for by Mrc1 as well as Chl1.

**Fig 6.**
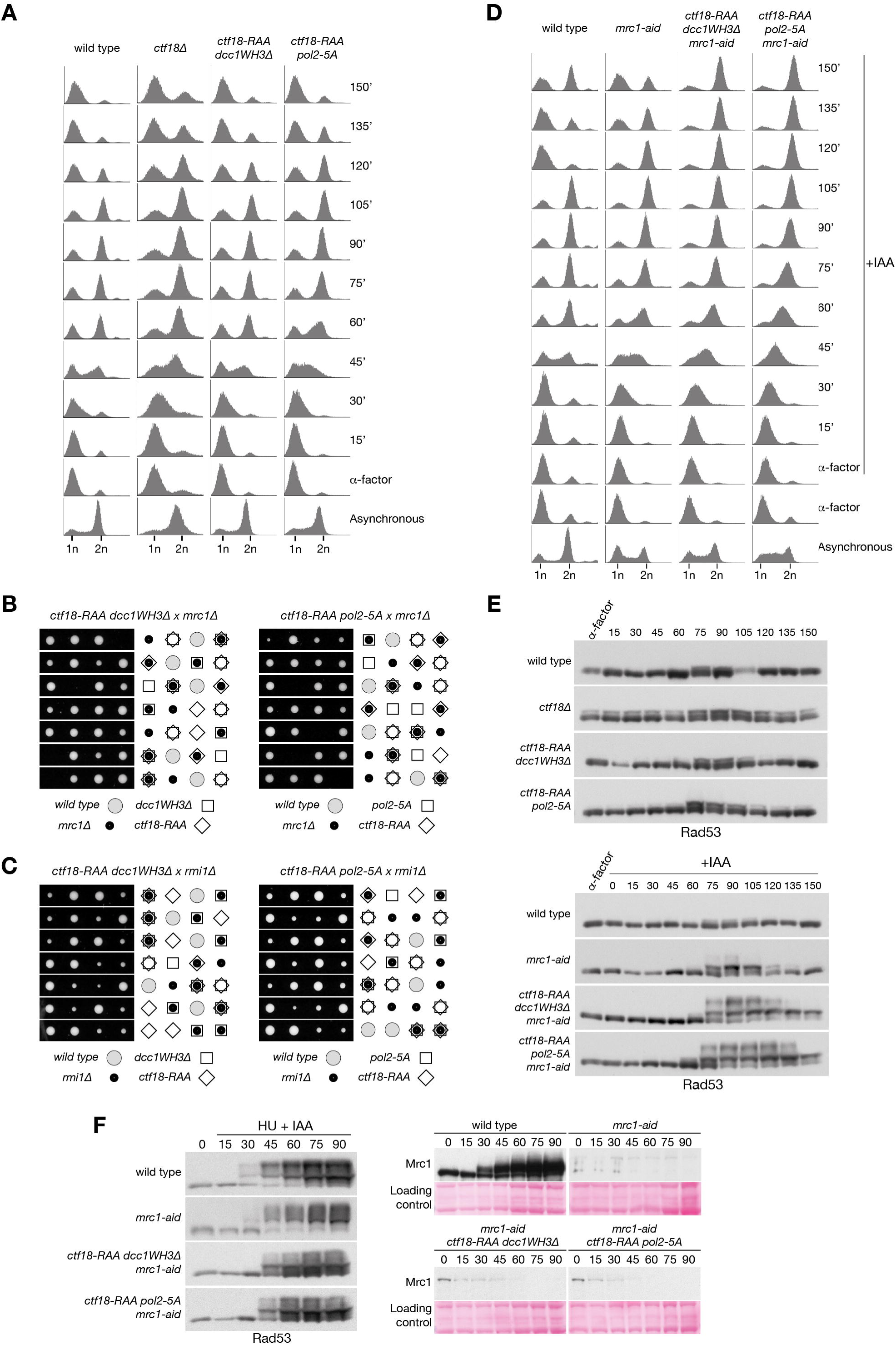
Mrc1 is essential for viability in the absence of the Ctf18-RFC/Pol ε interaction. **(A)** FACS analysis of DNA replication during a single cell cycle. The indicated strains were grown to exponential phase at 24°C, arrested in G1 and synchronously released. α-factor was added back 30 minutes after release to block entry in the next cell cycle. **(B-C)** Tetrad analysis of the meiotic progeny the diploids strains indicated, showing synthetic lethality between *ctf18-RAA dcc1WH3Δ/pol2-5A* and *mrc1Δ* **(B)**, but not *rmi1Δ* **(C). (D)** FACS analysis of DNA replication during a single cell cycle. The strains were treated as for panel A, except for the incubation for 1 hour in G1 with 0.5 mM IAA final concentration to induce the degradation of *mrc1-aid* and the release in medium containing 0.5 mM IAA. **(E)** Immunoblotting analysis of Rad53 phosphorylation from the experiments shown in panels **(A**,**D**). (F) Analysis of checkpoint activation following replication stress. Wild type, *mrc1-aid, mrc1-aid ctf18-RAA dcc1WH3Δ/pol2-5A* cells were arrested in G1, incubated with 0.5 mM IAA final concentration for 1 hour and released in medium containing 0.2 M HU and 0.5 mM IAA. Cells samples were collected at the indicated times and analyzed by immunoblotting for Rad53 (left) and Mrc1/mrc1-aid (right). Ponceau staining is shown for loading comparison.

### CTF18-RFC binding to Pol ε is required for the activation of the S phase checkpoint

To further characterize the dynamics of DNA replication in the *ctf18-RAA dcc1WH3Δ/pol2-5A* mutants, we analysed their levels of spontaneous recombination. Cells carrying a GFP-tagged version of Rad52 were arrested in G1 and synchronously released in S phase, either in the presence or absence of HU. Interestingly, in the presence of HU, both the *ctf18-RAA dcc1WH3Δ/ pol2-5A* mutants showed a great increase in Rad52 foci, similar to that observed in *ctf18Δ* cells (Fig. 7A,B). Since checkpoint activation is directly linked to repression of recombination (Alabert, Bianco and Pasero, 2009), the increased levels of recombination observed in the double mutants suggest a possible defect in checkpoint activation, as described for *CTF18* (Kubota *et al*., 2011; Crabbé *et al*., 2010). We first analysed whether the firing of late origins, which are inhibited following replication stress in a manner dependent on Rad53, Mrc1 and Ctf18 (Crabbé *et al*., 2010; Lopez-Mosqueda *et al*., 2010; Zegerman and Diffley, 2010), was defective in *ctf18-RAA dcc1WH3Δ/pol2-5A* mutants. We therefore analysed the levels of Psf1 phosphorylation in the replisome, a marker for late origin firing in HU (De Piccoli *et al*., 2012). We observed that, while the single *ctf18-RAA* mutation leads to a partial increase in Psf1 phosphorylation, a *ctf18-RAA pol2-5A* mutant shows a robust increase in Psf1 phosphorylation (Fig. 7C), similar to that previously observed in *ctf18Δ* cells, thus suggesting a defect in S phase checkpoint activation (García-Rodríguez *et al*., 2015). Analysis of Rad53 phosphorylation, however, did not show a marked decrease in phosphorylation for the *ctf18-RAA dcc1WH3Δ/pol2-5A* mutants (Fig. 7D), nor did the sensitivity to HU reflect severe S phase checkpoint defects (Fig. 4D;7E). The mild phenotype observed could be a consequence of the activation of the DNA damage checkpoint, which is also activated following exposure to HU, although with slower kinetics, thus partially compensating for the defects in Rad53 phosphorylation (Alcasabas *et al*., 2001; Katou *et al*., 2003). In fact, combining the *ctf18-RAA dcc1WH3Δ/pol2-5A* mutants with the deletion of the DNA damage checkpoint mediators *RAD24* or *RAD9* (de la Torre-Ruiz, Green and Lowndes, 1998; Weinert, Kiser and Hartwell, 1994), shows a marked decrease in Rad53 phosphorylation and leads to hypersensitivity to replication stalling (Fig. 7D,E; S6A,B). In addition, even single mutations of *DCC1, CTF18* or *POL2* lead to Rad53 activation and HU sensitivity phenotypes in the absence of *RAD24*, suggesting that even mild perturbation of the Pol ε/Ctf18-RFC interaction results in defective activation of the S phase checkpoint (Fig. S6C-E). Together, these observations highlight the importance of Ctf18-RFC recruitment to the leading strand for the replication stress response.

**Fig 7.**
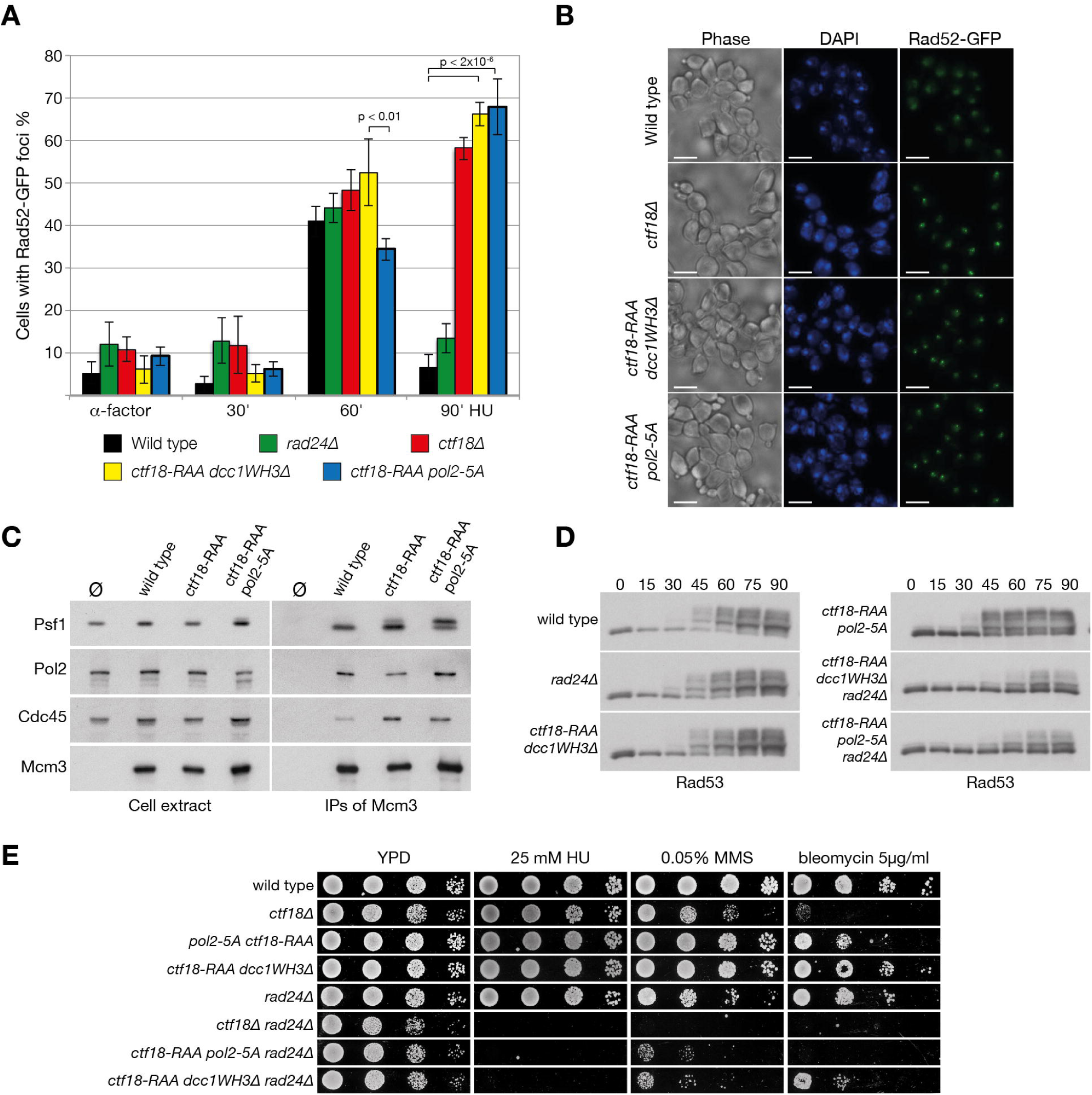
Loss of Ctf18-RFC from forks leads to defects in S phase checkpoint activation. **(A)** Analysis of the formation of Rad52-GFP foci in wild type, *ctf18Δ, rad24Δ, ctf18-RAA dcc1WH3Δ* and *ctf18-RAA pol2-5A* mutants. Cells were arrested in G1 and synchronously released in the presence or absence of 0.2 M HU, fixed, and analyzed by microscopy. The experiment was repeated at least three times and between 200-250 cells were counted for each sample. **(B)** Examples of the images from the analysis shown in *(A)*. The bar at the bottom of the images represent 5µm. **(C)** Disruption of the Pol2/Ctf18-1-8 interaction blocks the inhibition of late origin firing. Wild type, *ctf18-RAA and ctf18-RAA pol2-5* cells, all carrying a TAP-tagged allele of Mcm3, together with an untagged strain, were arrested in G1 and synchronously released in YPD 0.2 M HU for 90 min. Cells extracts were incubated with anti-TAP beads and the immunoprecipitated material was analyzed by immunoblotting. **(D)** Rad53 phosphorylation in response to replication fork stalling depends on the DNA damage checkpoint in cells defective in the Pol2-Ctf18-1-8 interaction. The indicated strains were arrested in G1 and synchronously released in medium containing 0.2 M HU. **(E)** Breaking the Pol ε/Ctf18-RFC interaction leads to synthetic replication stress sensitivity in the absence of *RAD24*. The strains were diluted 1:10 and spotted on the indicated medium.

## Discussion

Processive DNA replication, DNA repair, cohesion establishment and checkpoint activation are often intertwined; in fact, mutations or deletions of genes involved in DNA replication can affect several other processes as well. Understanding whether these phenotypes are the indirect consequence of large-scale defects in DNA replication and replisome conformation, or the direct results of specific protein binding and activation, requires dissecting the molecular mechanisms linked to these phenotypes. By analyzing how Ctf18-RFC is recruited to the leading strand via Pol ε, we have revealed that Ctf18-RFC has distinct functions in cohesion establishment, DNA replication and the activation of the S phase checkpoint.

### Pol2 recruits Ctf18-RFC to the leading strand

Our *in vitro* analysis demonstrates that the Ctf18-1-8 module of Ctf18-RFC forms a stable complex with the catalytic segment of Pol ε and that this interaction does not interfere with the catalytic activity or DNA binding of the polymerase, suggesting that the complex will be maintained throughout normal DNA replication. How does the large interface readily adapt to the conformational changes of the polymerase? We show this is achieved through an unusual protein-protein mechanism that utilizes electrostatic slippery interfaces to enable high binding plasticity. These low-specificity electrostatic patches explain why it was previously thought that the Ctf18-1-8 complex binds to DNA (Wade *et al*., 2017; Grabarczyk, Silkenat and Kisker, 2018).

Strikingly, we observe that, *in vivo*, the same complex forms at replication forks, both during DNA synthesis and fork stalling, as well as during the G1 phase of the cell cycle, confirming that the two complexes are constitutively associated (Fig 4A, S4E). Using our cryo-EM structure, we have designed mutations that specifically address the functions of the interaction between the two complexes while maintaining the integrity of the clamp loader. In agreement with the large binding surface, we observe an additive effect when mutating two different sites, resulting in the loss of Ctf18-RFC from forks, increased defects in checkpoint activation, replication stress sensitivity and synthetic interaction with several pathways in the cell (Fig S5D). Strikingly, the strength of the phenotype depends on the severity of the displacement of Ctf18-RFC from forks, closely linking the role of Ctf18-RFC to its localization by Pol ε.

One exception to this observation is that the loss of the Ctf18-RFC interaction with Pol ε and its displacement from the leading strand does not affect chromosome cohesion. Since *ctf18Δ* causes a loss of PCNA from forks (Lengronne *et al*., 2006) and its cohesion defects are can partially be suppressed by the deletion of *WPL1* and are epistatic with an *eco1Δ wpl1Δ* mutant allele (Borges *et al*., 2013), the role of Ctf18-RFC in chromosome cohesion appear to depend on its PCNA-loading activity and the resultant Eco1 recruitment and locking of cohesin onto the DNA (Moldovan, Pfander and Jentsch, 2006; Kenna and Skibbens, 2003; Rolef Ben-Shahar *et al*., 2008; Zhang *et al*., 2008; Unal *et al*., 2008). This suggests that *ctf18-RAA dcc1WH3Δ/pol2-5A* mutants load sufficient PCNA to support cohesion establishment. Whether this is mediated by recruitment of the mutant Ctf18-RFC near forks via alternative interactions or by some residual and transient binding to Pol2, which can only be removed by complete disruption of the Ctf18-1-8 module (Okimoto *et al*., 2016), is not understood.

### A stable Ctf18-RFC/Pol ε complex is essential for the rapid signaling of stalled forks

We have observed that the binding of Ctf18-RFC to Pol ε is essential for S phase checkpoint activation. Strikingly, even mutations that slightly weaken this binding show severe defects in activation of the replication checkpoint upon HU treatment (Fig. S6D). How is the interaction critical for the detection of nucleotide depletion induced stalling of Pol ε? Our data show that dissociation of Pol2_CAT_ from its template does not affect its interaction with Ctf18-1-8 and that Ctf18-1-8 has no direct regulatory role on the polymerase. Instead, the interaction appears to act solely to position the clamp loader module of Ctf18-RFC in proximity to the template regardless of whether Pol2_CAT_ is synthesizing DNA or disengaged (Fig. 8A-B). Dissociation of Pol2_CAT_ from the template is a prerequisite for binding of the template by the clamp loader portion of Ctf18-RFC, since it competes with Pol2_CAT_ for the same substrate, and it has been shown that it cannot load PCNA when Pol2_CAT_ is synthesizing DNA (Fig. 8A) (Fujisawa *et al*., 2017). We propose that when Pol2_CAT_ is not synthesizing DNA due to nucleotide depletion, Ctf18-RFC is positioned by Pol ε to compete with it to bind to the primed template, and could then signal the stalled state (Fig. 8B). Previous *in vitro* data show that under different conditions, Ctf18-RFC can load PCNA, unload it or form a trapped state with it (Bylund and Burgers, 2005). Future detailed structural and functional studies of the clamp loader portion of Ctf18-RFC will be necessary to understand how it acts on the free junction.

**Fig 8.**
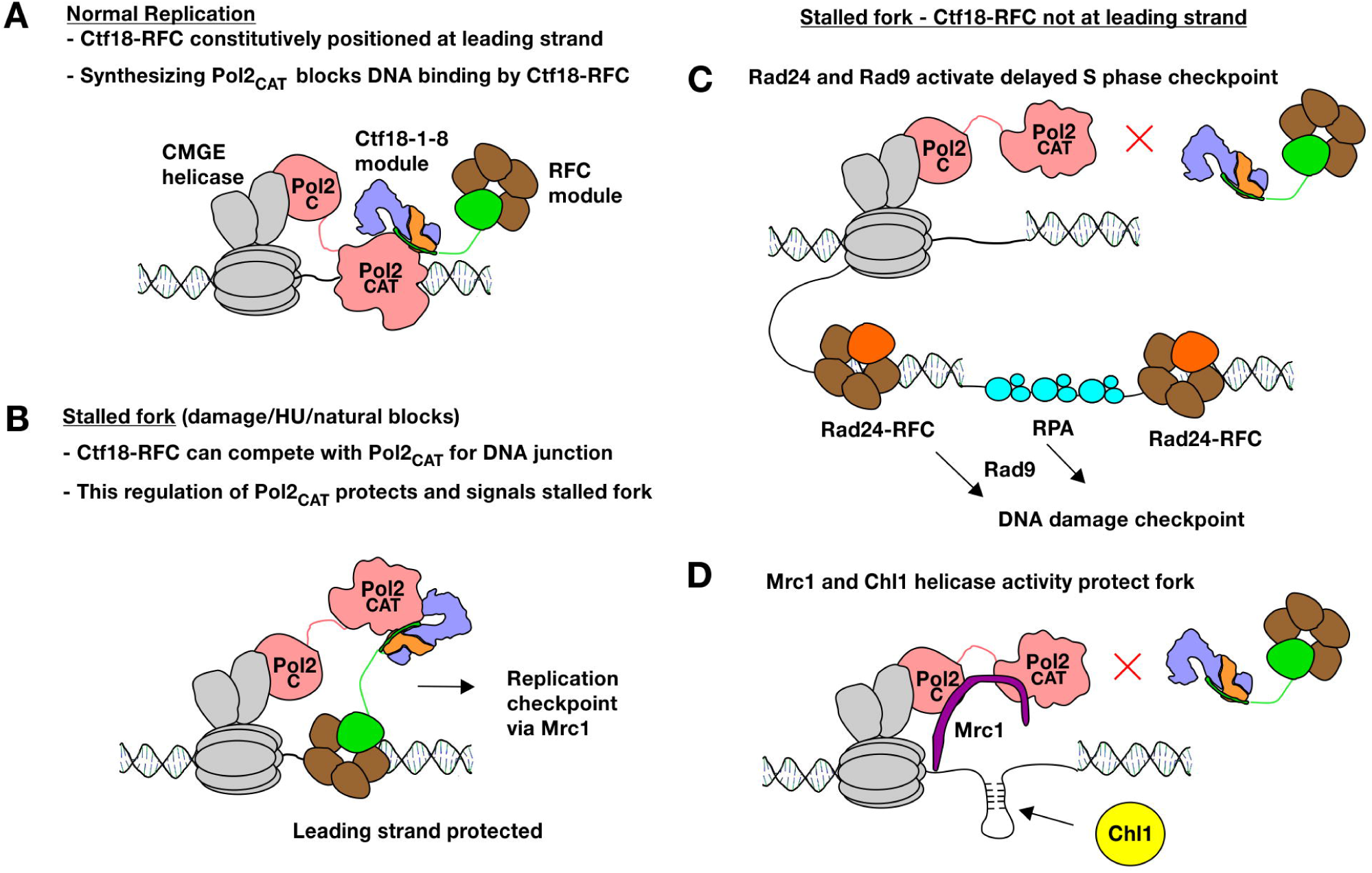
Model for the roles of Ctf18-RFC at the leading strand. **(A)** In wild-type cells, Ctf18-RFC is constitutively bound to Pol2_CAT_, and does not affect its activity. **(B)** When the leading strand stalls, Ctf18-RFC would be positioned to compete with Pol2_CAT_ for the DNA substrate. The positioning of Ctf18-RFC is essential for protecting and signaling the stalled fork. **(C)** The replication checkpoint requires Ctf18-RFC to be positioned at the leading strand. Absence of the replication checkpoint leads to gaps and overpriming which are signaled by the DNA damage checkpoint. **(D)** When Ctf18-RFC is not positioned at the leading strand the fork, it can no longer favor a protecting conformation of Pol2 resulting in replication defects. Mrc1 and the helicase activity of Chl1 compensate to protect the fork.

While downstream events are not clear at this stage, it is interesting to note that following phosphorylation by Mec1, the Mrc1 N-terminus no longer interacts with Pol2 and promotes the recruitment of Rad53 to the replisome, leading to its phosphorylation and activation (Lou *et al*., 2008; Osborn and Elledge, 2003). A possible explanation for this mechanism is that, by the process described above (Fig. 8B), Ctf18-RFC promotes the correct orientation of the Pol2_CAT_-Mrc1 complex at stalled forks in order to allow the timely phosphorylation of the Mrc1 N-terminus, thus initiating the activation of the S phase checkpoint. Direct detection of Pol ε stalling by Ctf18-RFC and Mrc1 would be consistent with the rapid kinetics of the replication checkpoint, compared to slower activation of the compensatory *RAD24* and *RAD9*-mediated DNA damage checkpoint by the slow build-up of unreplicated gaps and overpriming (Bacal *et al*., 2018) (Fig. 8C).

### A checkpoint-independent role in ensuring leading strand synthesis

Our genetic data point to a key role for the Pol ε/Ctf18-RFC interaction during DNA replication; in its absence, cell viability depends on the presence of Mrc1. However, this is not caused by defects in checkpoint activation (Fig. 6F), nor appears to depend on defects in chromosome cohesion (Fig 5A,B, 6C). Instead, in the *mrc1-aid ctf18-RAA pol2-5A/dcc1WH3Δ* strains, cells accumulate DNA damage following DNA replication. In fact, while the bulk of DNA replication in the mutants occurs with kinetics similar to the controls, cells activate the DNA damage checkpoint and arrest in G2/M phase, indicating that gaps behind the fork or double-stranded breaks might accumulate.

As Ctf18-1-8 does not directly regulate Pol2_CAT_ activity, and the clamp loader portion of Ctf18-RFC can only access the DNA template when Pol2_CAT_ has dissociated from it, this suggests that the function of Ctf18-RFC in normal replication must also relate to fork stalling or pausing, but by natural obstacles or stochastic dissociation of Pol2_CAT_ from the template. One possibility is that if Pol2_CAT_ dissociates from the template, PCNA is lost, and Ctf18-RFC must reload it to ensure efficient DNA synthesis (Fujisawa *et al*., 2017). Like Ctf18-RFC, Mrc1 directly interacts with Pol2 (Lou *et al*., 2008), and as well as its checkpoint function, promotes fork progression and restart following stalling (Hodgson, Calzada and Karim, 2007; Tourrière *et al*., 2005; Yeeles *et al*., 2017). This could mean that synthetic lethality results from Mrc1 promoting efficient synthesis in the absence of Ctf18-RFC-loaded PCNA. However, in this case it is surprising that bulk DNA replication kinetics are unaffected in the triple mutant strains (Fig. 6D).

Alternatively, Mrc1 and Ctf18-RFC may have complementary roles in allowing stalled/paused forks to restart damage-free (Fig. 8B,D). An essential function for leading strand Ctf18-RFC in protecting forks is further supported by the synthetic lethality of our interaction mutants with the catalytically dead Chl1 mutant. It is believed that the human Chl1 orthologue DDX11 requires its helicase activity to protect stalled forks (Calì *et al*., 2016). DDX11 is as a 5’-3’ helicase that requires a minimal ssDNA tail and can unwind flaps, forked structures, D-loops and G4 structures (Farina *et al*., 2008; Wu *et al*., 2012). This suggests that, when Ctf18-RFC is absent from the leading strand, unprotected endogenously stalled forks might form aberrant DNA structures that require unwinding by Chl1 (Fig. 8D).

Finally, we observed that the mechanism of interaction between Ctf18-RFC and Pol ε appears to be conserved in human cells. (Fig. 4G) Since roles for CTF18-RFC in DNA damage repair, fork speed regulation and chromosome cohesion have been described in human cells (Ogi *et al*., 2010; Terret *et al*., 2009), understanding the structural and functional impact of the presence of Ctf18 at forks could have important consequences for the understanding the origin, development and treatment of human diseases such as cancer.

## Conflict of interest

The authors declare that they have no conflict of interest.

## Acknowledgements

We would like to thank Erik Johansson, Karim Labib, Frank Uhlmann and Stephen Royle for strains, antibodies and plasmids. We are grateful to members of our labs for feedback, and to Caroline Kisker and Ben Berks for critical reading of the manuscript. GDP, KS, AW and the experimental cost were funded by Cancer Research UK Career Development Fellowship C44595/A16326. DBG was supported by DFG grant GR 5152/3-1. We would like to acknowledge the help of the Media Preparation Facility in The School of Life Sciences at the University of Warwick. We also give thanks to Dr Erick Martins Ratamero at the Computational Advanced Microscopy Development Unit for support in the analysis of microscopy data. We are grateful for support from beamline scientists at the EMBL-operated PETRA III beamline P13 at DESY. We would like to thank Bettina Böttcher, Tim Rasmussen and Vanessa Flegler for operation of the North Bavarian Cryo-EM Facility (DFG INST 93/903-1 FUGG) and for the collection of the cryo-EM dataset.

## Author contributions

KS, AW, BS, GDP and DBG performed the experiments; GDP and DBG planned the work and wrote the paper.

## Materials and Methods

### Molecular Biology

Exonuclease-deficient Pol2 (D290A/E292A) (Morrison *et al*., 1991) was used for *in vitro* experiments to simplify assays involving DNA. pETm14-POL2(1-1192)^exo^ was generated by restriction cloning by amplifying POL2 from the full-length Pol2 plasmid using primers 5’-CAAGAGGAGCTCGGGGACAAGTATATGATGTTTGGC-3’ and 5’-CAAGAGCTCGAGTTACTTAAACTTATCCTCCTTTGTAGCG-3’, mixing with stop-codon deleted pETm14, digesting with SacI and XhoI and ligating. The Ctf18 VRK mutation was generated by two sequential blunt-end mutagenesis reactions, first with primers 5’-CGCAGGAAAAATGTGACTTGGAATAACCTG-3’ and 5’-AGCGTTAGAGAACCCCTCATTGTATTTCAC-3’ to generate Ctf18 V730R, and then primers 5’-GCAAATGTGACTTGGAATAACCTGTGGG-3’ and 5’-CGCGCGAGCGTTAGAGAACCCCTC-3’ to generate Ctf18 V730R/R731A/K732A. POLE1 cDNA sequence obtained from TransOMIC Technologies was incorporated as a restriction fragment into pEGFP-C1 (KpnI + XbaI) and pEGFP-N1 (KpnI + AgeI). Mutations were generated by fusion PCR and inserted in linearized plasmids (SalI-SrfI) using NEBuilder® HiFi DNA Assembly (New England BioLabs). Generation of the plasmids used for Yeast-Two-Hybrids were generated as in (García-Rodríguez *et al*., 2015).

### DNA substrates

The P1P2 substrate (Hogg *et al*., 2014) was generated by mixing oligonucleotides P1, 5’-TAACCGCGTTC-3’, and P2, 5’-CTCTTGAACGCGGTTA-3’, heating to 95°C and slow cooling to 4°C. For gel based assays and cryo-EM, P1* with a 5’ Cy3 fluorescent label was used for the annealing reaction. All oligonucleotides were purchased from Sigma-Aldrich.

### Protein purification

Ctf18-1-8 and Pol2(1-528) expression and purification were performed exactly as described previously (Grabarczyk, Silkenat and Kisker, 2018). For Pol2_CAT_, pETm14-POL2(1-1192)^exo^ was transformed into BL21(DE3)Star (Novagen) cells and grown in media supplemented with 50 µg/ml kanamycin. Large terrific broth expression cultures were inoculated 1 in 100 with an overnight start culture and grown at 30°C with shaking at 200 rpm until they reached an A_600_ of 0.5. The temperature was reduced to 17°C, and cultures were induced with 0.4 mM IPTG overnight, followed by harvesting using centrifugation, and storage of bacterial pellets at −80°C until use. Thawed pellets were resuspended in 50 mM Tris-HCl pH 8.0, 500 mM NaCl, 1 mM TCEP, 10 mM imidazole, 1 EDTA-free protease inhibitor tablet/50 mL (Roche), and 30 U/mL DNase I. Lysis was performed by two passages through a Microfluidics M-110P microfluidizer at 150 MPa. The lysate was cleared by centrifugation for 1 hour at 60000 x g and then loaded on a 5 mL Histrap FF column (GE Healthcare) equilibrated in 50 mM Tris-HCl pH 8.0, 500 mM NaCl, 1 mM TCEP, using an Akta Purifier FPLC system. The column was then washed with 16 column volumes of 25 mM imidazole in the same buffer, and then eluted with a 10 column volume gradient of 25 – 250 mM imidazole. Protein-containing fractions were then concentrated by microfiltration before loading on a Superdex 200 26/60 prepgrade column (GE Healthcare) column that had been equilibrated in 20 mM HEPES pH 7.5, 160 mM NaCl, 1 mM TCEP. Pol2_CAT_ containing fractions were diluted 1 in 4 into 40 mM Tris pH 8.0, 160 mM NaCl, 1 mM TCEP, and then loaded on a MonoQ 10/100 GL column (GE Healthcare) equilibrated in 25 mM Tris pH 8.0, 160 mM NaCl, 1 mM TCEP. The protein was eluted with a 180 to 580 mM NaCl gradient over 18 column volumes. Fractions containing Pol2CAT were then concentrated and buffer-exchanged into 20 mM HEPES pH 7.5, 160 mM NaCl, 1 mM TCEP using a spin concentrator and stored at −80°C. The protein concentration was initially estimated by the A_280_ absorption using a calculated extinction coefficient ε_280_ of 153450 M^-1^cm^-1^.

### Crystallization

Pol2(1-528) and Ctf18-1-8 were mixed at a 1:1 ratio and purified by size-exclusion chromatography using a Superdex 200 10/300 gl column (GE Healthcare) as described previously. Crystallization trials were performed with a Honeybee fluid transfer robot using the sitting drop vapor-diffusion method with 0.3 µL of protein mixed with 0.3 µL of mother liquor. Drops were incubated at 20°C and equilibrated against 40 µL mother liquor. The Pol2(1-528)/Ctf18-1-8 complex was crystallized at 5 mg/ml and mixed with a precipitant solution containing 0.2 M NaCl, 14% PEG 20000, 0.1 M HEPES pH 7.0. Crystals were cryo-protected in the same solution with 25% ethylene glycol additional.

### X-ray data collection, structure solution and refinement

X-ray data were collected at 1.0332 Å at DESY beamline P13. The data were integrated, scaled and merged using XDS (Kabsch, 2010) followed by Aimless (Evans and Murshudov, 2013). The structure was solved by sequential molecular replacement using Phaser (McCoy *et al*., 2007) and fragments of pdb 5oki as a model. First, we searched for two copies of Pol2(1-528) with all possible screw axes. This was successful with a TFZ score of 15.9. This revealed the correct space group as P3_2_21. Extra density was visible which corresponded approximately to the position of Dcc1 WH2-3 in the C222_1_ crystal form, and so these were manually docked into both copies and rigid fitted in Coot (Emsley *et al*., 2010) and then refined using Phenix Refine (Adams *et al*., 2010). A new round of molecular replacement was performed searching for this entire model and one copy of Ctf18-1-8 TBD-WHL-WH1. Following this, the second copy of Ctf18-1-8 TBD-WHL-WH1 was then docked using non-crystallographic symmetry. After linking the fragmented structure, the model was refined using Phenix Refine (Adams *et al*., 2010), with reference restraints from the high resolution structures, secondary structure restraints, NCS restraints and group B factors. Coot (Emsley *et al*., 2010) was used to rigid body fit loops and delete parts not present in the density. Refinement resulted in R_work_/R_free_ values of 27.5/33.3% with 95.0% Ramanchandran favored residues, and 0.35% outliers.

### Cryo-EM sample and grid preparation and data collection

Samples were prepared both in the presence and absence of the P1P2* substrate and ultimately resulted in the same structure, but only the sample with DNA is discussed as the grids were of much higher quality. 2 µM Ctf18-1-8, Pol2CAT and P1P2* in buffer 20 mM HEPES pH 7.5, 160 mM NaCl, 1 mM TCEP were incubated on ice for half an hour and then applied on top of a 4 mL 10 to 30% w/v sucrose density gradient containing 0.15% v/v EM grade glutaraldehyde (EMS). The density gradient was prepared using a Biocomp Gradient Master™ according to the Grafix steps (Stark, 2010). This was then centrifuged in a SW60-Ti rotor for 18 hours at 45000 rpm and 4°C. Fractions were removed with a pipette and analyzed by SDS-PAGE. Fractions containing a ∼200 kDa complex with no additional aggregate were first examined by negative stain EM for particle homogeneity. The best fraction was desalted into buffer containing 20 mM HEPES pH 7.5, 100 mM NaCl, 1 mM TCEP to remove sucrose. 4 µL of sample (∼0.1 mg/ml) was applied to a glow-discharged R 1.2/1.3 Cu 400 Quantifoil grid, blotted for 3 s, force 10 under 100% humidity and vitrified by plunging into liquid ethane using a VitroBot Mark IV (Thermo Fisher). Data were collected using a Thermo Fisher Titan Krios G3 microscope operating at 300kV with a Falcon IIIEC detector in counting mode. The nominal magnification was 75000 with calibrated pixel size of 1.0635 Å /pix. Defocus values range from −1.4 to −2.6 µm. 2205 micrographs were collected, each comprising of a 47-frame movie with 75s exposure and accumulated dose of 60 e/Å^2^.

### Cryo-EM image processing and model building

Motion correction and dose-weighting was performed on the fly during data collection using MotionCor2 (Zheng *et al*., 2017). Initial processing of the summed micrographs was performed in cisTEM (Grant, Rohou and Grigorieff, 2018). Gaussian blob picked particles were subjected to 2D classification and the best classes used to reconstruct an ab initio 3D model which showed clear secondary structure. However, the initial map could not be improved without overfitting noise. Therefore, we continued processing with Relion 3.0 beta (Zivanov *et al*., 2018). After CTF estimation with CTFFIND-4.1 (Rohou and Grigorieff, 2015), ∼1000 particles were manually picked and subjected to reference-free 2D classification. Three classes with approximately the correct dimensions were chosen as templates for auto-picking, which found 1240576 particles. These were extracted with a box size of 290 pixels and downscaled to 64 pixels. All following steps except the final refinement were performed with a 150 Å circular mask. Three rounds of reference-free 2D classification were used to remove ice and other contaminants, resulting in 903024 protein particles. 3D classification was performed using the initial model from cisTEM lowpass-filtered to 60 Å as a reference. The 311977 particles from good classes were reextracted in their original pixel size and could be refined to 4.5 Å resolution after masking. They were then subjected to another round of 3D classification into 5 classes, with the angular sampling and then local angular sampling increased successively throughout classification. One class had no particles. The best class (‘Class 1’) was refined and reached 4.3 Å resolution after masking, and was re-refined using this mask to increase alignment accuracy. The final resolution as judged by an FSC of 0.143 was 4.2 Å. Sharpening was performed with a fitted negative B-factor of −143 Å^2^ and the map was filtered to 4.2 Å. Local resolution analysis was performed with ResMap (Kucukelbir, Sigworth and Tagare, 2013). To obtain the second class, the best two classes from the second round of classification were subjected to a third round of 3D classification with alignment into 8 classes. Two of the classes strongly resembled ‘Class 1’, while two others has a distinct positioning of Ctf18-1-8. The best of these (‘Class 2’) was subjected to a single refinement using a circular mask, and after masking had a resolution of 5.8 Å judged by an FSC of 0.143. The map was sharpened with a B factor of −156 Å2 and filtered to 5.8 Å. Modelling and refinement were performed using the sharpened maps. Models from a combination of pdb 4m8o, pdb 5okc and pdb 5oki were docked into the highest resolution map using UCSF Chimera (Pettersen et al., 2004), and subjected to multiple rounds of building and refinement using Coot (Emsley et al., 2010) and Phenix Real Space Refine (Adams et al., 2010). The 4Fe4S cluster was modelled using pdb 6h1v (Ter Beek *et al*., 2019). Secondary structure and reference model restraints (from pdb 4m8o and 5okc) were used throughout. For Class 2, the model from Class 1 was docked into the density using UCSF Chimera (Pettersen *et al*., 2004) and subjected to 25 cycles of refinement with Phenix Real Space Refine (Adams et al., 2010). Structural figures were made using either the PyMol Molecular Graphics System or UCSF Chimera (Pettersen et al., 2004).

### Isothermal titration calorimetry

ITC experiments were performed at 25°C using a MicroCal iTC_200_ in 20 mM HEPES pH 7.5, 160 mM NaCl, 1 mM TCEP, with a stirring speed of 1000 rpm and a reference power of 11 cal/sec. Control experiments were performed to ensure that sample dilution did not cause systematic deviation from a flat baseline in the presence and absence of 50 µM P1P2. Integrated values obtained with OriginPro (OriginLab) were fit to a 1:1 hetero-association model using SEDPHAT (Houtman *et al*., 2007). Errors in the protein concentration determination were accounted for by including a value for the incompetent fraction of Pol2_CAT_ based on the clear mid-point of the titration. Titrations were replicated with separate protein preparations.

### Gel-based polymerase activity and DNA binding assays

All gels containing Cy3-labelled DNA were scanned with a Pharos FX fluorescence imaging system (Biorad) using excitation/emission wavelengths of 532/605 nm. All *in vitro* gel-based experiments were replicated with a separate biological protein preparation. The concentration of Pol2_CAT_ was corrected for the inactive fraction observed in ITC. Polymerase primer extension assays were performed in buffer 25 mM HEPES pH 7.5, 60 mM KCl, 8 mM MgOAc, 0.1 mM dNTPs, 0.1 mg/ml BSA, 10% glycerol, 1 mM TCEP and used 100 nM substrate P1P2*. The reaction was initiated by adding the indicated concentration of Pol2_CAT_ or a Pol2_CAT_/Ctf18-1-8 mixture, and incubated at room temperature for 5 minutes. The reaction was quenched by 1:1 addition of SDS, EDTA, and the sample was denatured by incubation at 99°C for two minutes and immediate transfer to ice for one minute. The samples were immediately loaded on a 7M urea Tris-acetate-EDTA 15% polyacrylamide gel and run at 70 V at room temperature and then imaged. EMSAs were performed in buffer 25 mM HEPES pH 7.5, 60 mM KCl, 10% glycerol, 1 mM TCEP and used 50 nM substrate P1P2*. The indicated protein concentrations were added and the samples incubated on ice for 20 minutes. Samples were then loaded on a pre-run 6% Tris-glycine polyacrylamide gel and run at 70 V at 4°C and then imaged.

### Yeast strains and cell growth

Budding yeast *Saccharomyces cerevisiae* strains used are in the Yeast Strains List. All strains were derived from W303-1a, containing the alleles: *ade2-1, ura3-1, his3-11,15 trp1-1, leu2-3,112, rad5*-535, *can1-100*. Yeast was grown in YP medium supplemented with either glucose (YPD), galactose (YPGAL) or raffinose (YPRAF) to a final concentration of 2% (w/v). When grown on solid media, a final agar concentration of 1% (w/v) was used. Temperatures of 24, 30, 35 and 37°C were used for growth, based on viability or experimental design. For cell cycle experiments, cultures were diluted from an initial inoculum to give a final cell density of 0.3-0.4×10^7^ cells/ml in the required volume. Cells were then grown until they reached a cell density of 0.7-0.8×10^7^ cells/ml. To arrest cells in G1 α-factor was added at 7.5μg/ml final concentration, after 90 minutes α-factor was added every 30 minutes at 3.25μg/ml final concentration to maintain cells in G1.

In experiments utilizing galactose-inducible protein expression, cells were inoculated overnight in YPRAF and arrested in G1 in YPRAF for 3 hours, before being switched to YPGAL for 40 minutes in the presence of α-factor. To release from α-factor arrest, cells were washed 2 times using fresh medium lacking α-factor. Cultures were then resuspended in YPD or in medium containing 0.2M HU. For protein samples analysis, 10 ml cultures were collected, resuspended in 20% TCA, and treated as in (De Piccoli *et al*., 2012).

For cell spotting experiments, strains were streaked from glycerol storage suspensions at −80°C onto YPD agar plates and incubated at 24°C until colonies were sufficiently large. Four to six colonies were then re-suspended in 1mL of sterile water and diluted to create concentrations of 0.5×10^6^, 0.5×10^5^, 0.5×10^4^ and 0.5×10^3^ cells/ml in sterile water. 10μL of each dilution was spotted in line onto square plates of YPD and YPD containing selective compounds HU, MMS, Bleomycin or Benomyl. Plates were then left to grow at the desired temperature for up to five days and scanned every 24 hours.

Hela cells were grown in DMEM medium supplemented 10% FBS at 37°C in humid atmosphere with 5% CO2. POLE1 variants fused to GFP were transiently transfected using GeneJuice Transfection Reagent (Novagene, cat. no. 70967-5) according to the manufacturer protocol. Cells were harvested 48 hours after transfection and processed into IP.

### Harvesting yeast cells for IP

Yeast cultures were washed with cold 4°C 20mM HEPES-KOH pH7.9, followed by a wash with 100mM HEPES-KOH ph7.9, 100mM KOAc, 10mM MgOAc, 2mM EDTA, again at 4°C. After washing the pellet was suspended in a fresh volume of the same solution, supplemented with 2 mM glycerophosphate (Johnson Matthey), 2 mM NaF (Fisher), 1 mM DTT, 1% (v/v) Sigma protease inhibitor cocktail (for fungal and yeast extracts, Sigma) and 0.24% (w/v) EDTA-free Complete Protease Inhibitor Cocktail (Roche). Ratio of cell pellet mass to final mass of suspension was either 1:3 (250mL culture) or 4:1 (1L culture). For 1L cultures, inhibitor concentration was increased to 8 mM glycerophosphate, 8 mM NaF, 1 mM DTT, 4% (v/v) Sigma protease inhibitor cocktail (for fungal and yeast 78 extracts) and 0.48% (w/v) EDTA-free Complete Protease Inhibitor Cocktail. Suspensions were snap-frozen by pipetting drop-wise into liquid nitrogen.

For cross-linking experiments, formaldehyde was added to the culture to a final concentration of 1%. Samples were cross-linked for fifteen minutes unless otherwise stated. Glycine was added to a final concentration of 1.3 M and incubated for five minutes to arrest cross-linking. Cells to be harvested were washed with cold 20 mM HEPES-KOH pH 7.9, followed by a wash with cold 50 mM HEPES-KOH pH 7.9, 140 mM NaCl, 1 mM EDTA. The pellet was then suspended in a volume of the same solution (ratio 1:3) supplemented with 2 mM glycerophosphate (Johnson Matthey), 2 mM NaF (Fisher), 1% (v/v) Sigma protease inhibitor cocktail (for fungal and yeast extracts, Sigma) and 0.24% (w/v) EDTA-free Complete Protease Inhibitor Cocktail (Roche).

### Protein immunoprecipitation and protein analysis

Cell extracts were generated using a 6870 FreezerMill (SPEX SamplePrep). To 1 g of thawed lysate (taken to be equivalent to 1 ml) a ¼ volume of 50% (v/v) glycerol, 100 mM HEPES-KOH (pH 7.9), 100 mM KOAc, 50 mM MgOAc, 0.5% Igepal CA-630 (Sigma), 2 mM EDTA supplemented with 2 mM glycerophosphate, 2 mM NaF, 1 mM DTT, 1% (v/v) Sigma protease inhibitor cocktail and 0.24% (w/v) EDTA-free Complete Protease Inhibitor Cocktail buffer was added. This solution was then incubated for 30 minutes with PierceTM Universal Nuclease for Cell Lysis (ThermoFisher) at 0.4 units/μl (for 250 mL culture samples) or 1.6 units/μl (for 1L culture samples) concentration at 4°C on a rotating wheel. Samples were then centrifuged step-wise at 18,700 g and then 126,600 g. The remaining supernatant was incubated with antibody-conjugated beads for 2 hours. Beads were then washed three times with a solution of 100 mM HEPES-KOH (pH 7.9), 100 mM KOAc, 50 mM MgOAc, 2 mM EDTA, 0.1% (v/v) Igepal® CA-630, once with 2 mM glycerophosphate, 2 mM NaF, 1 mM DTT, 1% (v/v) Sigma protease inhibitor cocktail and 0.24% (w/v) EDTA-free Complete Protease Inhibitor Cocktail. Samples were eluted by boiling with Lemmli buffer.

For cross-linked samples, samples were treated as in (De Piccoli *et al*., 2012). Briefly, the thawed lysate was mixed with a volume ¼ of it’s weight of 50 mM Hepes-KOH pH 7.5, 140 mM NaCl, 5% TritonX-100, 0.5% Sodium deoxycholate, 1 mM EDTA supplemented with 2 mM glycerophosphate, 2 mM NaF, 1% (v/v) Sigma protease inhibitor cocktail and 0.24% (w/v) EDTA-free Complete Protease Inhibitor Cocktail. Samples were then split into 500 μL aliquots and sonicated using a Soniprep 150 Plus sonication probe for seven cycles of fifteen seconds at power setting five. Samples were then spun in a microfuge at 13200rpm for fifteen minutes at 4°C. Supernatants were incubated for two hours at 4°C with antibody-conjugated beads.

Following incubation, beads were washed twice in 50 mM Hepes-KOH pH 7.5, 140 mM NaCl, 1% TritonX-100, 0.1% Sodium deoxycholate, 1 mM EDTA supplemented with 2 mM glycerophosphate, 2 mM NaF, 1% (v/v) Sigma protease inhibitor cocktail and 0.24% (w/v) EDTA-free Complete Protease Inhibitor Cocktail, twice in 50 mM Hepes-KOH pH 7.5, 500 mM NaCl, 1% TritonX-100, 0.1% Sodium deoxycholate, 1 mM EDTA supplemented with 2 mM glycerophosphate, 2 mM NaF, 1% (v/v) Sigma protease inhibitor cocktail and 0.24% (w/v) EDTA-free Complete Protease Inhibitor Cocktail and twice in 10 mM Tris-HCl pH 8.0, 250 mM LiCl, 0.5% NP-40, 0.5% Sodium deoxycholate, 1 mM EDTA. Finally, beads were washed once with 1 mL of TE, resuspended in 50 mM Tris-HCl pH 8.0, 10 mM EDTA, 1% SDS and incubated at 65°C for ten minutes. Following this, the supernatant and cell extracts were boiled for thirty minutes with Laemmli buffer.

Mass spectrometry analysis was conducted with the Proteomics Research Technology Platform at the University of Warwick.

### GFP-IP protocol

HeLa cells harvested after 48 h since transient transfection were washed twice in cold PBS and resuspended in 1 mL of cold lysis buffer (50 mM HEPES-KOH pH 7.5, 100 mM KAc, 1mM MgOAc, 0.1% Igepal, 10% glycerol) supplemented with 100 U/ml of Pierce Universal Nuclease (Thermo Scientific, cat no. 88702), 2x Protease inhibitors (Roche, cat no. 04693159001) and 1 mM DTT. Cells were lysed for 1 h on a rotation wheel at 4°C and spun for 20 min at 14000 x g at 4°C. Supernatant was collected in a fresh tube and 50 ul of each input sample was put aside and mixed with Laemmli buffer. 25 µl of GFP TrapA slurry beads (Chromotek) were added to the rest of the protein extracts and incubated for another 2 h on a rotation wheel in the cold room. Then samples were gently spun and beads underwent 3 washes in the washing buffer (lysis buffer without nuclease). Proteins bound to the beads were washed out with 2x Laemmli buffer, each for 5 min at 75°C with shaking. Samples were then run for WB.

### Western Blots

Protein samples were run on polyacrylamide gels (5-12%) and transferred onto nitrocellulose membrane. The bands were then probed with the appropriate primary antibody for 1 hour in 5% (w/v) milk in TBS-T. Membranes were then washed three times for 5 minutes in fresh TBS-T, incubated with the appropriate secondary antibody and washed again three times. The membranes were then treated with Western blotting reagents and the chemiluminescent signal was captured using films. Protein analysis of Pol epsilon and Ctf18-RFC was performed twice by technical repeat.

### Flow cytometry

Samples were collected and prepared as in (De Piccoli *et al*., 2012). Cells were analyzed using an LSRII flow cytometer and Cytoflow software.

### Microscopy

All experiments were independently conducted three times (biological repeats); samples for mutants and wild type controls strains were collected during the same experiment. Samples for microscopy were incubated for ten minutes on a rotating wheel at 24°C with methanol-free formaldehyde (final concentration of 8%). Samples were washed with 1 mL PBS and stored in PBS in the dark at 4°C for up to 24 hours. Samples were incubated with 1 μg/ml DAPI/PBS, imaged using a DeltaVision 2 microscope (Applied Precision Instruments). Images were analyzed using ImageJ and the number of foci were counted manually and, for the experiments shown in Fig 5B, using a macro (Sup Methods 1). The macro was validated by comparison with manual counting.

### DATA AND SOFTWARE AVAILABILITY

The X-ray structural model and structure factors reported in this study have been deposited in the Protein Data Bank with accession code 6S1C. Cryo-EM maps have been deposited in the Electron Microscopy Data Bank with accession codes 10088 and 10089, and the corresponding models in the Protein Data Bank with accession codes 6S2E and 6S2F. All other data supporting the findings of this study are contained within the published article and the supplementary material.

## YEAST STRAINS LIST

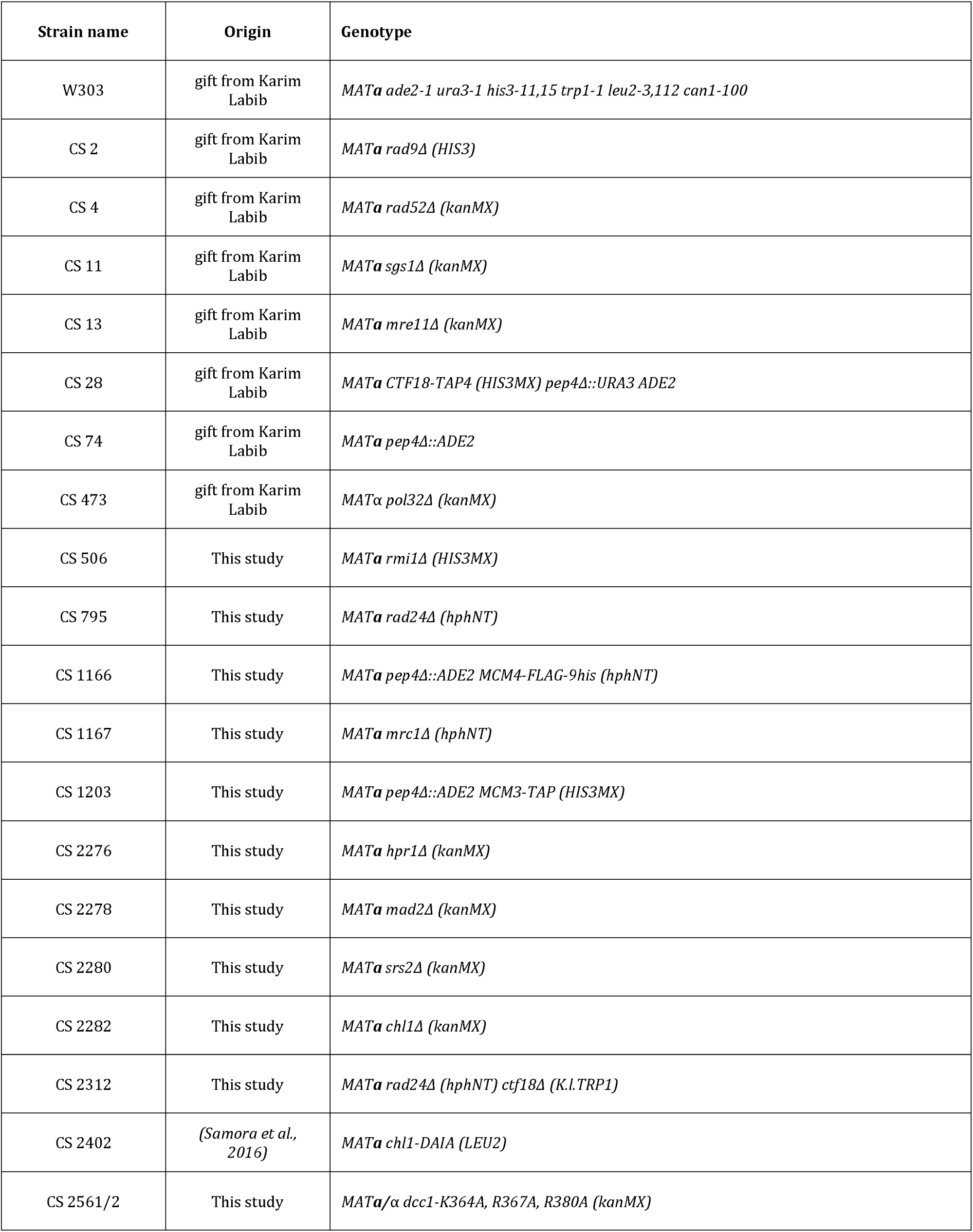
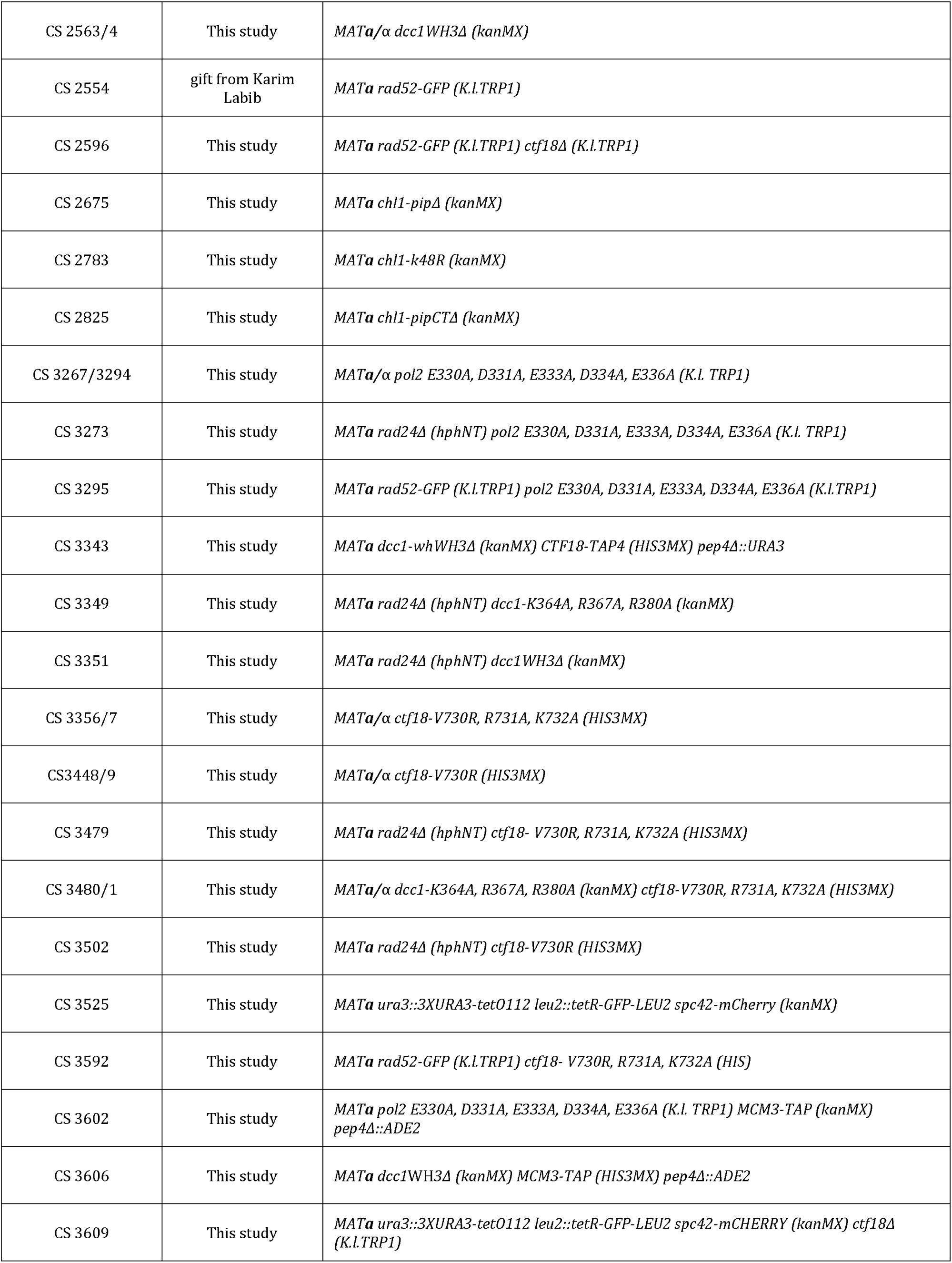
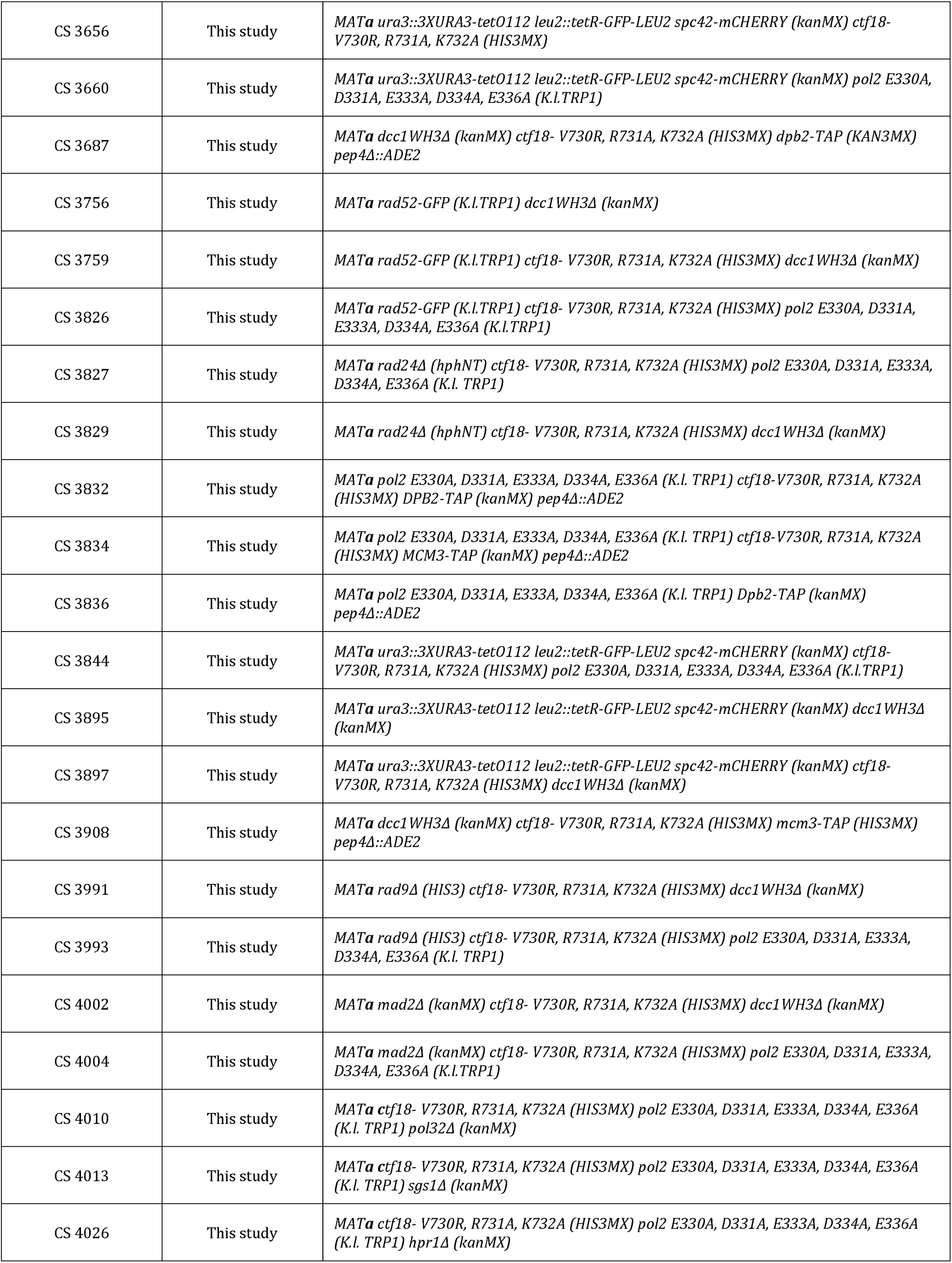
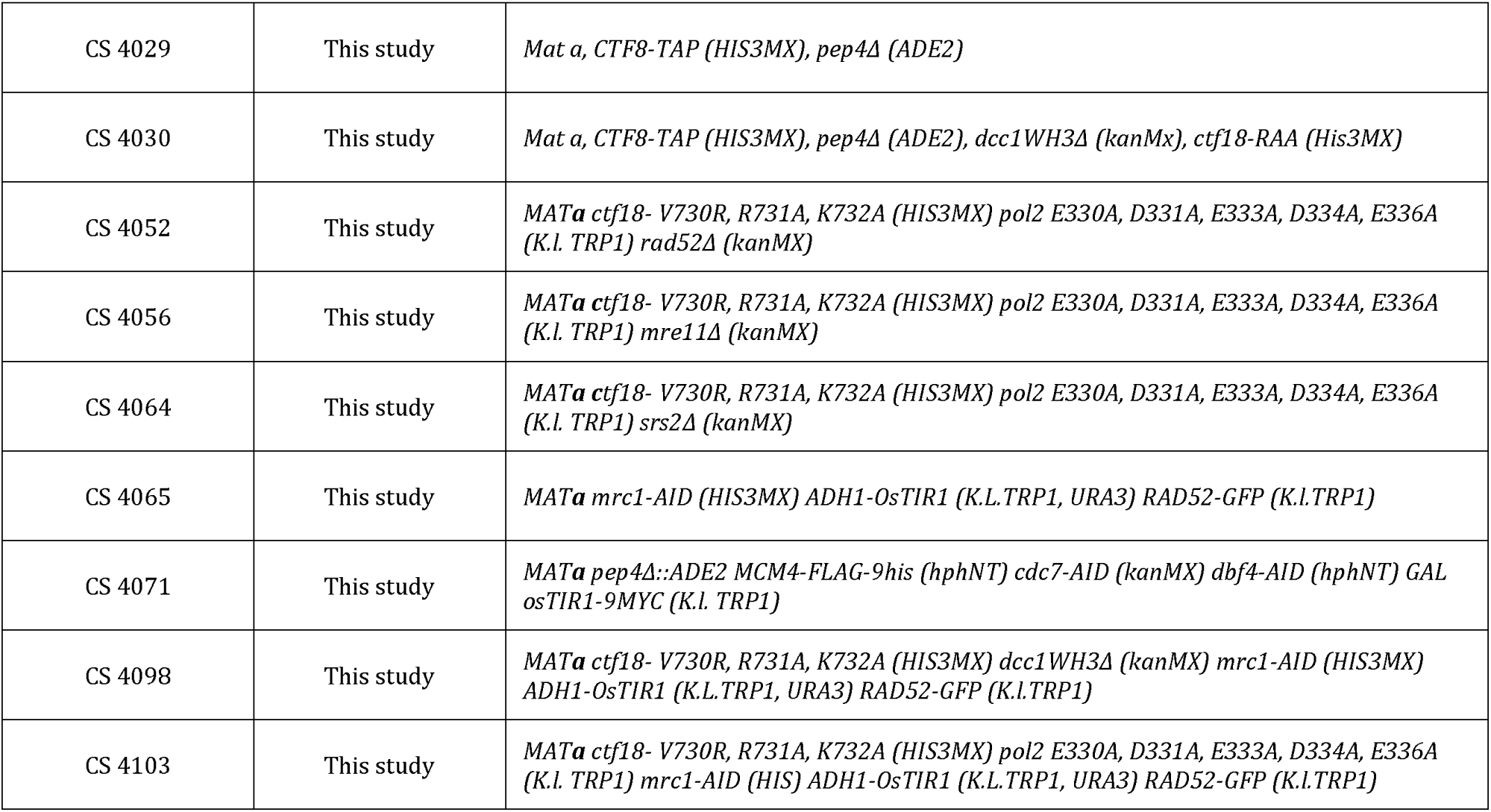

| **Name** | **Plasmids** |
| --- | --- |
| pCS62 | pGBKT7-Pol2 NT (1-396) mutant A |
| pCS63 | pGBKT7-Pol2 NT (1-396) mutant B |
| pCS64 | pGBKT7-Pol2 NT (1-396) mutant C |
| pCS65 | pGBKT7-Pol2 NT (1-396) mutant D |
| pCS66 | pGBKT7-Pol2 NT (1-396) mutant E |
| pCS67 | pGBKT7-Pol2 NT (1-396) mutant F |
| pCS68 | pGBKT7-Pol2 NT (1-396) mutant G |
| pCS69 | pGBKT7-Pol2 NT (1-396) mutant H |
| pCS70 | pGBKT7-Pol2 NT (1-396) mutant I |
| pCS71 | pGBKT7-Pol2 NT (1-396) mutant L |
| pCS85 | pEGFP-C1 POLE1 m134 |
| pCS86 | pEGFP-C1 POLE1 wt |
| pCS88 | pEGFP-N1 POLE1 m134 |
| pCS91 | EGFP-N1 POLE1 wt |
| pCS160 | pEGFP-C1 acidic |
| pCS162 | pEGFP-N1 acidic |
| pCS94 | pGADT7-CHTF18 wild type |
| pCS97 | pGBKT7-POLE1 1-450 |
| pGDP33 | pGBKT7-POLE1 1-450 m134 |
| pKL559 | pGADT7 |
| pKL560 | pGBKT7 |
| pLG1 | pGADT7-CTF18 wild type (García-Rodríguez *et al.*, 2015) |
| pLG55 | pGBKT7-Pol2 NT (1-396) wild type (García-Rodríguez *et al.*, 2015) |

| **Chemical and critical reagents** | **Manufacturer** |
| --- | --- |
| HU | Molekula LTD (10872383 \| 127-07-1) |
| MMS | Sigma-Aldrich (129925) |
| Bleomycin | VWR ( 203401) |
| Benomyl (PESTANAL (R) , analytical standard) | Sigma-Aldrich (45339) |
| Yeast extract (Bacto™) | Appleton Woods |
| Peptone (Oxoid) | Fisher Scientific (10404013) |
| Yeast nitrogen base (Difco™) | Appleton Woods |
| Alpha factor | Pepceuticals |
| Sigma protease inhibitor cocktail | Sigma-Aldrich (P8215) |
| EDTA-free Complete Protease Inhibitor Cocktail | Roche (11873580001) |
| Glycerophosphate | ALFA (AESAR) (L03425.22) |
| Dynabeads (M-270 Epoxy) | Invitrogen |
| Pierce™ Universal Nuclease for Cell Lysis | ThermoFisher (88700) |
| Amersham ECL Western blotting reagents | GE Healthcare (RPN 2209) |
| Anti-mouse-HRP | New England Biolabs (UK) LTD (7076S) |
| Anti-sheep-HRP | Sigma (S1265) |
| Anti-rabbit-HRP | Cell Signaling (7074S) |
| Anti-FLAG M2 | Sigma (F1804) |
| Anti-Dcc1 | gift from Karim Labib |
| Anti-Pol2 | gift from Karim Labib |
| Anti-Pol1 | gift from Karim Labib |
| Anti-Cdc45 | gift from Karim Labib |
| Anti-Ctf4 | gift from Karim Labib |
| Anti-Psf1 | gift from Karim Labib |
| Anti-Rad53 | Abcam PLC [EL7.E1] (ab166859) |
| Anti-POLE1 | Abcam PLC (ab226848) |
| Anti-CHTF18 | Abcam PLC (ab220288) |

