## Supplementary Information for "A stable leading strand polymerase/clamp loader complex is required for normal and perturbed eukaryotic DNA replication"

### Supplemental Figure Legends

**Fig S1. Structural and biophysical characterization of the Ctf18-1-8/Pol2$_{CAT}$ interaction. (A)** Superposition of the WH2 and WH3 domains of the C222$_1$ Ctf18-1-8/Pol2(1-528) crystal form (pdb 4oki), with Ctf8 shown in orange, Dcc1 in blue, Ctf18 in green and Pol2(1-528) in gray, and the new P3$_2$21 crystal form shown entirely in salmon. (**B**) Surface conservation calculated using the Consurf server of Pol2$_{CAT}$ and Ctf18-1-8 with the interacting surface indicated by the dashed yellow lines. Areas with least conservation are shown in cyan, and most conservation in magenta. (**C**) Comparison of subdomain movements of Pol2$_{CAT}$ (our structure and pdb 4m8o) with *E. coli* Pol II (pdb 3k5o and 3k59) (Wang and Yang, 2009) Structures are aligned by the NTD and EXO subdomains while subdomains indicated above are shown in red for the empty polymerase and salmon for the DNA and nucleotide-bound structure. (**D**) ITC titrations of Ctf18-1-8 in to Pol2$_{CAT}$ in the presence and absence of excess DNA substrate. (**E**) Comparison of the molecular details of the Dcc1-Pol2$_{EXO}$ interface for the Pol2$_{CAT}$/Ctf18-1-8 complex (left panel) and Pol2(1-528) complex (right panel).

**Fig S2. Cryo-EM data processing strategy.**

**Fig S3. Fit of cryo-EM models in maps.** (**A**) Refined model and cryo-EM electron density for Class 1. (**B**) Refined model and cryo-EM electron density for Class 2. (**C**) Sidechain electron density of a well-ordered region of Class 1. (**D**) Electron density surrounding the iron-sulfur cluster of Pol2$_{CAT}$.

**Fig S4. Breaking the interaction between Pol2 and Ctf18-RFC. A**) Analysis of the interaction between Pol ε and Ctf18-RFC during the cell cycle. The experiment was conducted as in Fig. 4A, except the cultures were not treated with formaldehyde. (**B**) Alignment of the N-terminal sequences of POL2 and orthologues in in *Schizosaccharomyces pombe*, Homo sapiens, *Mus musculus*, *Drosophila melanogaster* and *Xenopus laevis*. The sites selected for mutation and the correspondent amino acids substitution used in the screening are indicated. (**C**) Yeast-two hybrids analysis of the interaction between alleles of Pol2 (1-396) and Ctf18 (full-length). Cells were co-transformed with the plasmids as indicated. (**D**) Ctf18-RFC binding to Mcm2-7 depends on origin firing. Strains carrying auxin-inducible degrons (Nishimura *et al.*, 2009) *cdc7-aid dbf4-aid* and an *MCM4-5FLAG* allele, together with a *CDC7 DBF4 Mcm4-5FLAG* and untagged control strains, were grown in YPRAF to exponential phase at 24˚C, arrested in G1, resuspended in YPGAL for 35 minutes and incubated a further hour in the presence of 0.5 mM Indole-3-acetic acid for 1 hour. Cells were then synchronously released in S phase for 30 minutes and treated with formaldehyde. Cells extracts and the proteins immunoprecipitated with anti-FLAG beads were analyzed by immunoblotting. (**E**) The binding of Ctf18-RFC at forks does not increase following fork stalling. Strains, carrying a TAP-tagged allele of MCM3 or an untagged control, were arrested in G1, and synchronously released in YPD for 30 minutes (S phase) or YPD 0.2 M HU for 90 minutes (HU). Cells were treated with formaldehyde; the cells extracts and the proteins immunoprecipitated with anti-TAP beads were analyzed by immunoblotting. (**F**) Yeast-two hybrids analysis of the interaction between alleles of Pole1 (1-450) and Chtf18 (full-length). Cells were co-transformed with the plasmids as indicated. 3-amino-1,2,4-triazole (3-AT) was added to the final concentration of 0.75 mM to increase the specificity of the yeast two hybrids interaction.

**Fig S5 Analysis of the genetic interactions of the Pol2 and Ctf18-1-8 mutants.** (**A**) (Top). Graphical representation of Chl1, showing the position the conserved helicase motifs (from I to VI, in red), K48 (mutated in the catalytic-dead mutant), the Fe-S cluster (Fe-S), the Arch domain (Arch), the Ctf4-Interacting-Peptide (CIP, 700-712) and the putative PCNA-Interacting-Peptide (PIP, 833-840) (Cortone *et al.*, 2018). (Bottom). Deletion of the PIP motif in *CHL1* causes DNA damage sensitivity similar to that observed in the catalytic-dead allele chl1K48R. The indicated strains were diluted 1:10 and spotted on the specified medium. (**B**) *chl1PIPΔ* is lethal in a *ctf18-RAA pol2-5A* background. Analysis of the meiotic progeny of diploids heterozygotes for *chl1PIPΔ ctf18-RAA* and *pol2-5A* is shown. Plates were incubated at 24˚C and scanned after 3 days growth. (**C**) Analysis of the meiotic progeny of the indicated heterozygotes diploids. No synthetic lethality was observed, but growth defects with *ctf18-RAA pol2-5A* were observed in combination *sgs1Δ, mre11Δ, pol32Δ and srs2Δ*. Plates were incubated at 24˚C and scanned after 3 days. (**D**) Analysis of the synthetic defects. The indicated strains were diluted 1:10 and spotted on the specified medium.

**Fig S6. Checkpoint activation in response to fork stalling largely depends on the DNA damage checkpoint in mutants defective in the Pol ε/Ctf18-RFC interaction.** (**A**) Analysis of checkpoint activation following replication stress. Wild type, *rad9Δ*, *rad9Δ ctf18-RAA dcc1WH3/pol2-5A* cells were arrested in G1 and synchronously released in medium containing 0.2 M HU and 0.5 mM IAA. Cells samples were collected at the indicated times and analyzed by immunoblotting for Rad53. (**B**)

Deletion of *RAD9* is confers synthetic sensitivity to HU in *ctf18-RAA dcc1WH3/pol2-5A* strains. The strains were diluted 1:10 and spotted on the indicated medium. (**C**) Analysis of the dynamics of Rad53 phosphorylation in Pol2/Ctf18-RFC complex single mutants, in the presence or absence of *RAD24*. The indicated strains were treated as in panel A. (**D**) Analysis of HU sensitivity in Pol2/Ctf18-RFC complex single mutants, in the presence or absence of *RAD24*. The strains were diluted 1:10 and spotted on the indicated medium. (**E**) Analysis of double mutants for Ctf18 and Dcc1 show synthetic defects in HU sensitivity. The strains were diluted 1:10 and spotted on the indicated medium.

**Table S1. Cryo-EM data collection and refinement statistics**

**Table S2. X-ray data collection and refinement statistics**

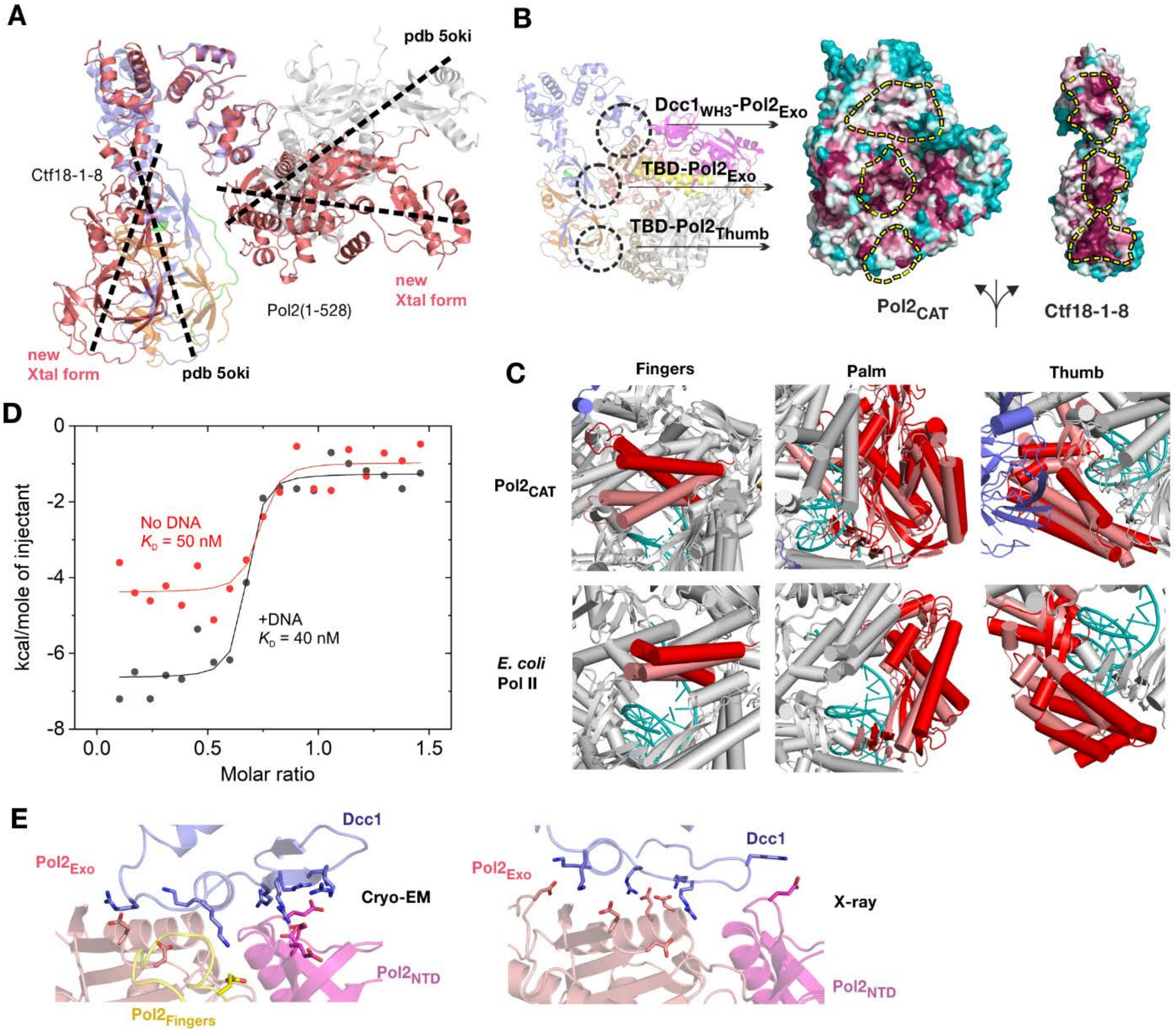

A
pdb 5oki
Ctf18-1-8
new Xtal form
Pol2(1-528)
new Xtal form
pdb 5oki
B
Dcc1WH3-Pol2Exo
TBD-Pol2Exo
TBD-Pol2Thumb
Pol2CAT
Ctf18-1-8
C
Fingers
Palm
Thumb
Pol2CAT
E. coli Pol II
D
No DNA
KD = 50 nM
+DNA
KD = 40 nM
kcal/mole of injectant
Molar ratio
E
Pol2Exo
Dcc1
Cryo-EM
Pol2NTD
Pol2Fingers
Pol2Exo
Dcc1
X-ray
Pol2NTD


**Fig. S1**

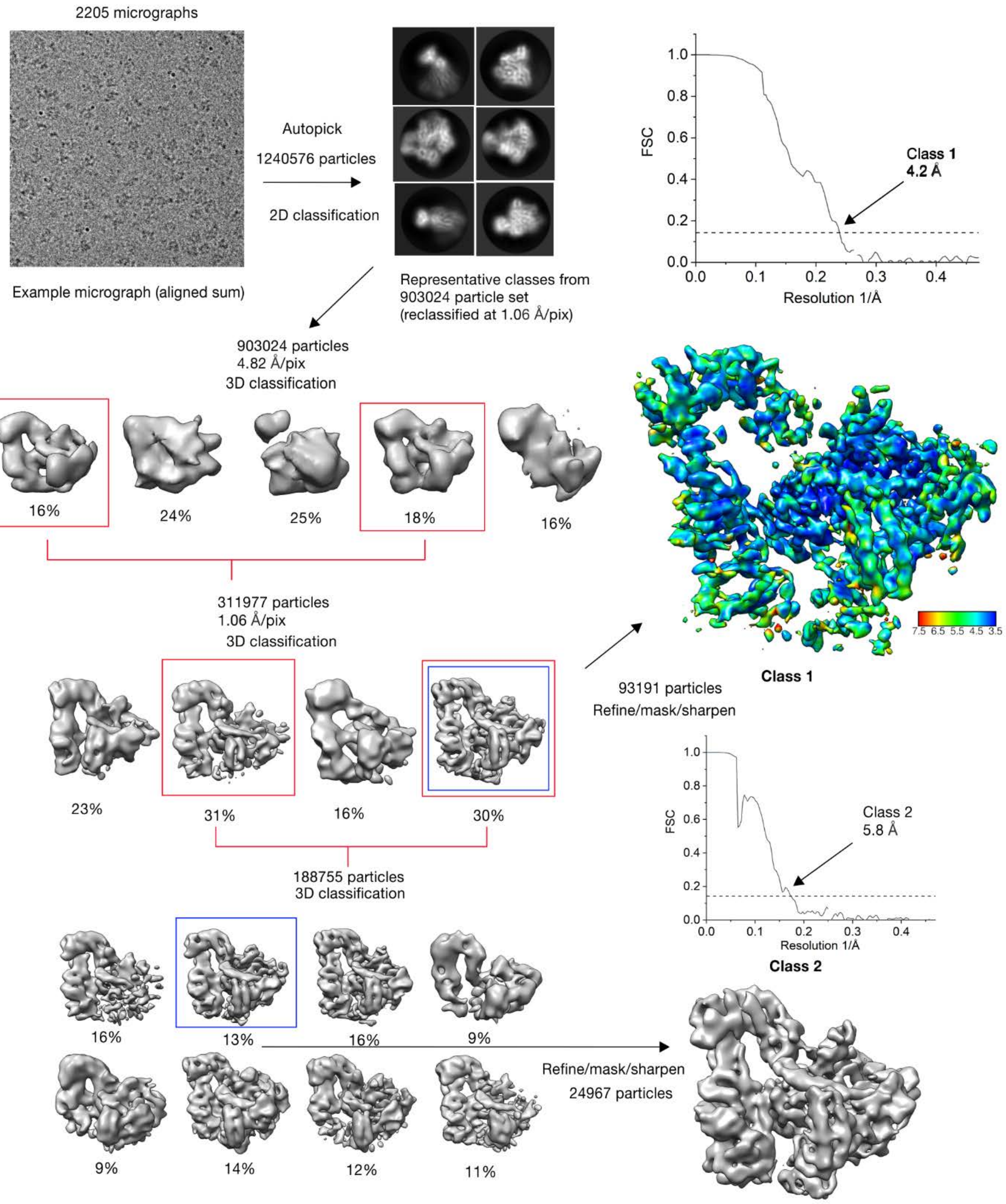
2205 micrographs
Example micrograph (aligned sum)
Autopick
1240576 particles
2D classification
Representative classes from
903024 particle set
(reclassified at 1.06 Å/pix)
FSC
Resolution 1/Å
Class 1
4.2 Å
903024 particles
4.82 Å/pix
3D classification
16%
24%
25%
18%
16%
311977 particles
1.06 Å/pix
3D classification
23%
31%
16%
30%
188755 particles
3D classification
7.5 6.5 5.5 4.5 3.5
Class 1
93191 particles
Refine/mask/sharpen
Class 2
5.8 Å
Class 2
16%
13%
16%
9%
9%
14%
12%
11%
Refine/mask/sharpen
24967 particles

**Fig S2**

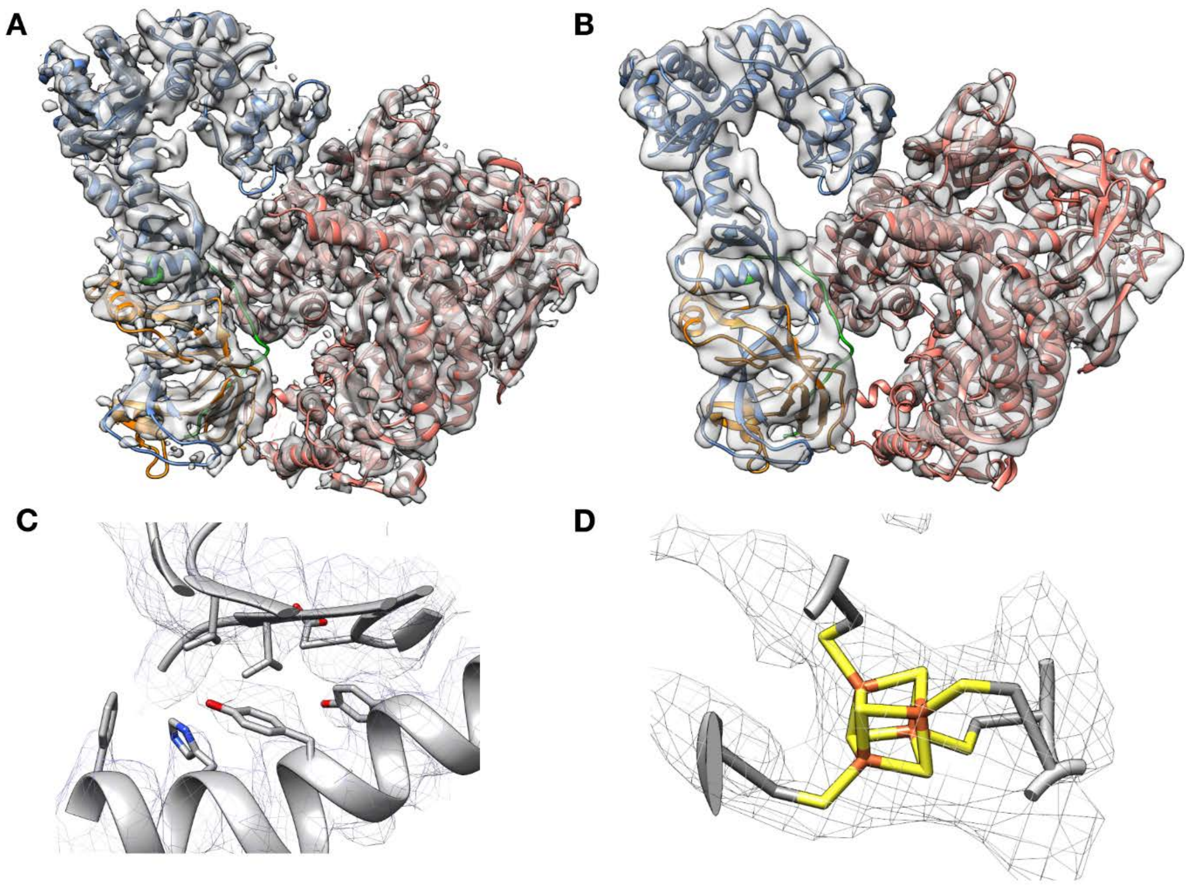


**Fig S3**

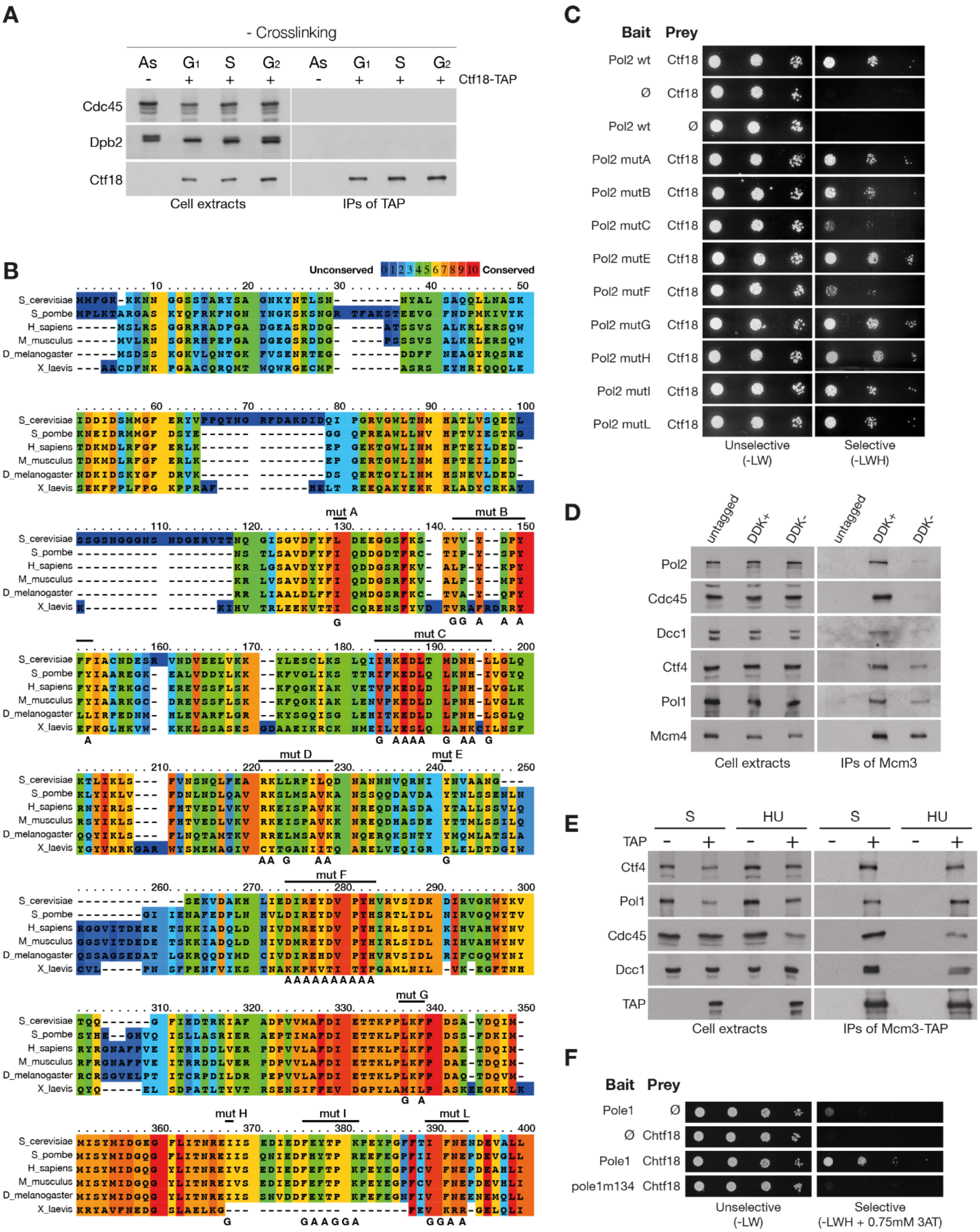
A
- Crosslinking
As G1 S G2 As G1 S G2
Ctf18-TAP
Cdc45
Dpb2
Ctf18
Cell extracts
IPs of TAP
B
Unconserved
Conserved
S_cerevisiae
S_pombe
H_sapiens
M_musculus
D_melanogaster
X_laevis
mut A
mut B
mut C
mut D
mut E
mut F
mut G
mut H
mut I
mut L
C
Bait
Prey
Pol2 wt
Ctf18
Ø
Pol2 mutA
Pol2 mutB
Pol2 mutC
Pol2 mutE
Pol2 mutF
Pol2 mutG
Pol2 mutH
Pol2 mutI
Pol2 mutL
Unselective
(-LW)
Selective
(-LWH)
D
untagged
DDK+
DDK-
Pol2
Cdc45
Dcc1
Ctf4
Pol1
Mcm4
Cell extracts
IPs of Mcm3
E
S
HU
TAP
Ctf4
Pol1
Cdc45
Dcc1
TAP
Cell extracts
IPs of Mcm3-TAP
F
Bait
Prey
Pole1
Ø
Chtf18
pole1m134
Unselective
(-LW)
Selective
(-LWH + 0.75mM 3AT)

**Figure S4**

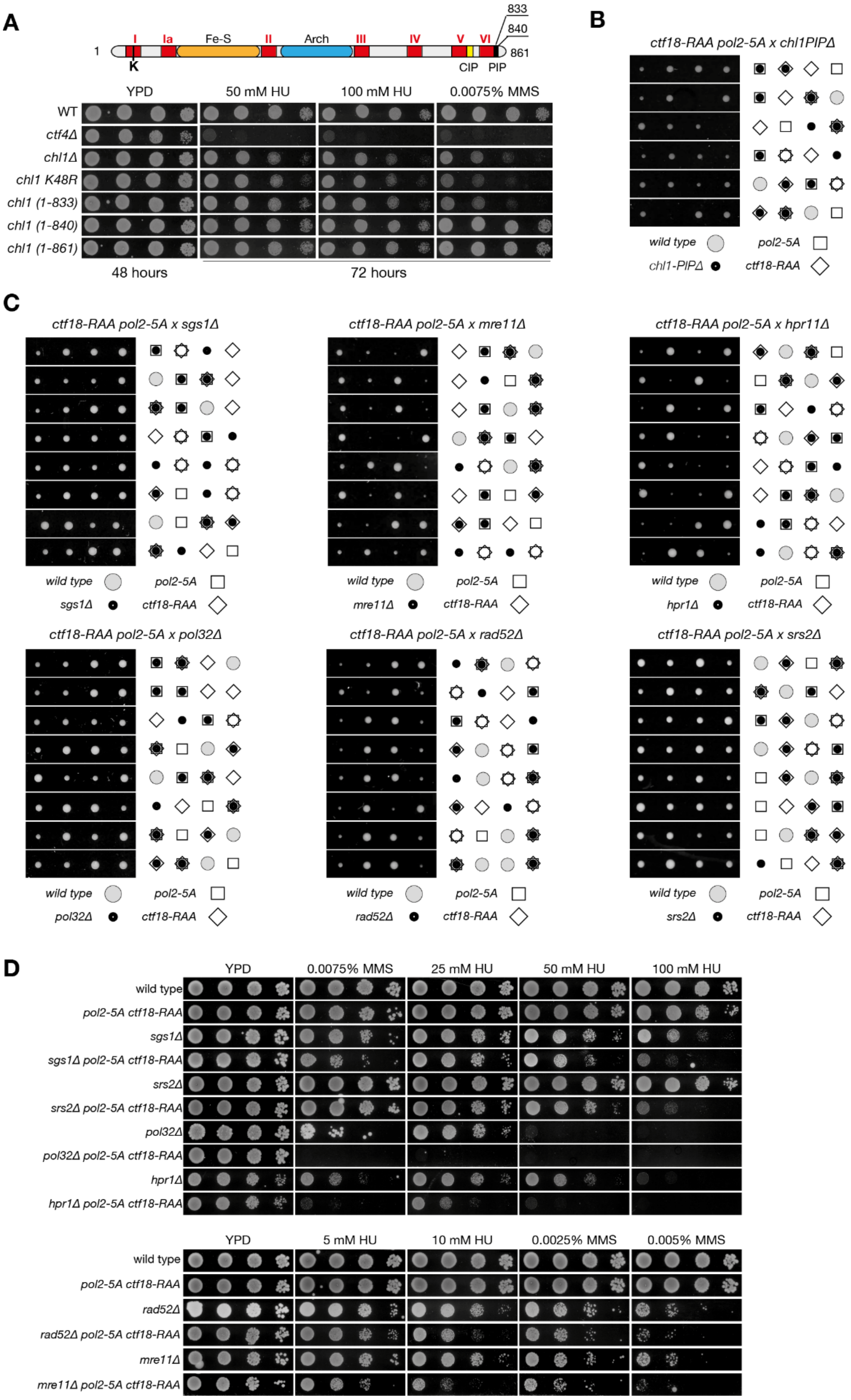
A
1
I
Ia
Fe-S
II
Arch
III
IV
V
VI
833
840
861
K
CIP
PIP
YPD
50 mM HU
100 mM HU
0.0075% MMS
WT
ctf4Δ
chl1Δ
chl1 K48R
chl1 (1-833)
chl1 (1-840)
chl1 (1-861)
48 hours
72 hours
B
ctf18-RAA pol2-5A x chl1PIPΔ
wild type
pol2-5A
chl1-PIPΔ
ctf18-RAA
C
ctf18-RAA pol2-5A x sgs1Δ
ctf18-RAA pol2-5A x mre11Δ
ctf18-RAA pol2-5A x hpr11Δ
wild type
pol2-5A
sgs1Δ
ctf18-RAA
mre11Δ
hpr1Δ
ctf18-RAA pol2-5A x pol32Δ
ctf18-RAA pol2-5A x rad52Δ
ctf18-RAA pol2-5A x srs2Δ
pol32Δ
rad52Δ
srs2Δ
D
YPD
0.0075% MMS
25 mM HU
50 mM HU
100 mM HU
wild type
pol2-5A ctf18-RAA
sgs1Δ
sgs1Δ pol2-5A ctf18-RAA
srs2Δ
srs2Δ pol2-5A ctf18-RAA
pol32Δ
pol32Δ pol2-5A ctf18-RAA
hpr1Δ
hpr1Δ pol2-5A ctf18-RAA
YPD
5 mM HU
10 mM HU
0.0025% MMS
0.005% MMS
wild type
pol2-5A ctf18-RAA
rad52Δ
rad52Δ pol2-5A ctf18-RAA
mre11Δ
mre11Δ pol2-5A ctf18-RAA

**Figure S5**

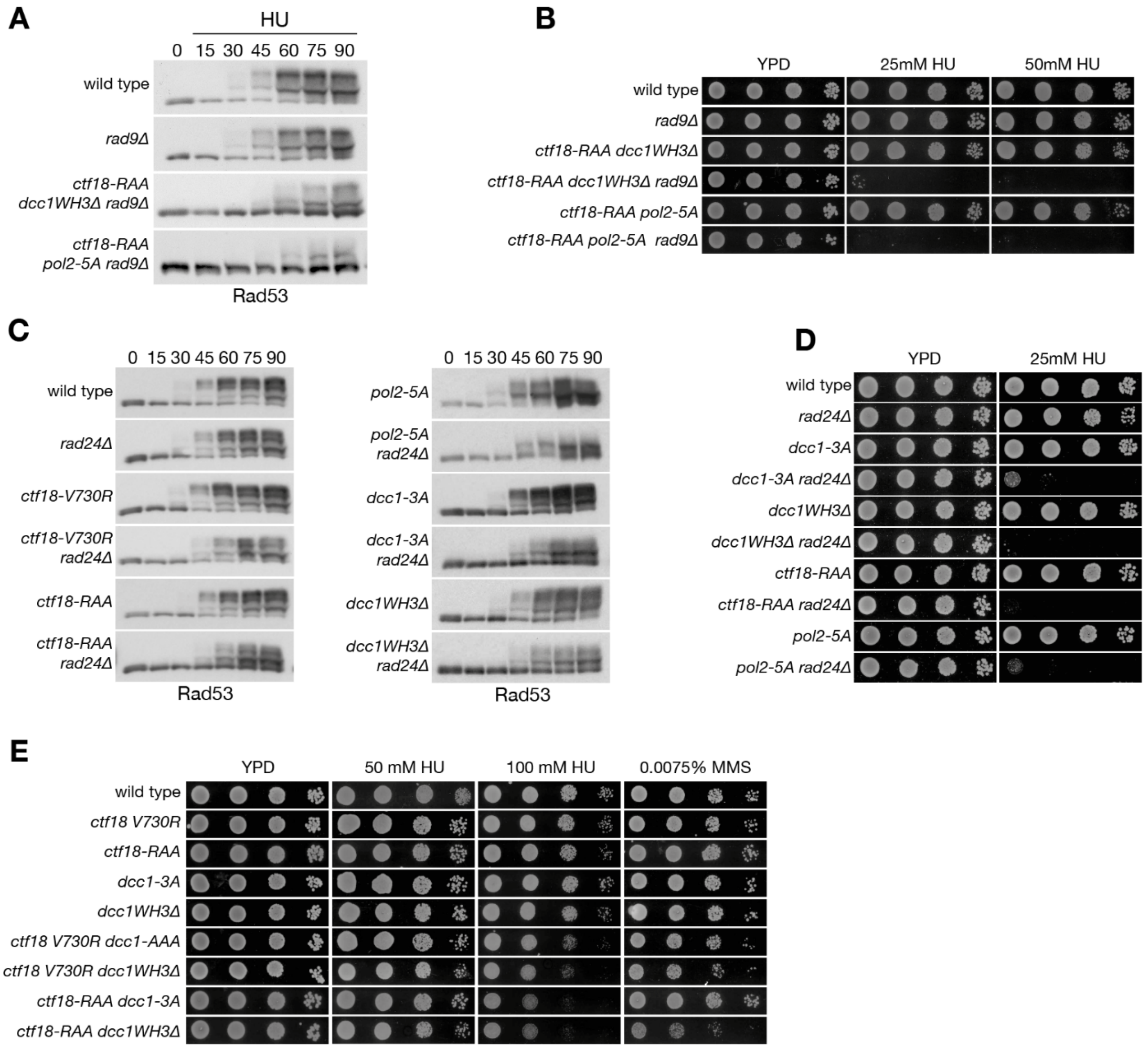

A
HU
0 15 30 45 60 75 90
wild type
rad9Δ
ctf18-RAA dcc1WH3Δ rad9Δ
ctf18-RAA pol2-5A rad9Δ
Rad53
B
YPD
25mM HU
50mM HU
wild type
rad9Δ
ctf18-RAA dcc1WH3Δ
ctf18-RAA dcc1WH3Δ rad9Δ
ctf18-RAA pol2-5A
ctf18-RAA pol2-5A rad9Δ
C
0 15 30 45 60 75 90
wild type
rad24Δ
ctf18-V730R
ctf18-V730R rad24Δ
ctf18-RAA
ctf18-RAA rad24Δ
Rad53
pol2-5A
pol2-5A rad24Δ
dcc1-3A
dcc1-3A rad24Δ
dcc1WH3Δ
dcc1WH3Δ rad24Δ
Rad53
D
YPD
25mM HU
wild type
rad24Δ
dcc1-3A
dcc1-3A rad24Δ
dcc1WH3Δ
dcc1WH3Δ rad24Δ
ctf18-RAA
ctf18-RAA rad24Δ
pol2-5A
pol2-5A rad24Δ
E
YPD
50 mM HU
100 mM HU
0.0075% MMS
wild type
ctf18 V730R
ctf18-RAA
dcc1-3A
dcc1WH3Δ
ctf18 V730R dcc1-AAA
ctf18 V730R dcc1WH3Δ
ctf18-RAA dcc1-3A
ctf18-RAA dcc1WH3Δ


**Figure S6**

**Table S1. Cryo-EM data collection and refinement statistics**

| | $Pol2_{CAT}$-Ctf18-1-8 Class 1 | $Pol2_{CAT}$-Ctf18-1-8 Class 2 |
|---|---|---|
| **Data collection and processing** | | |
| Magnification | 75,000 | 75,000 |
| Voltage (kV) | 300 | 300 |
| Electron exposure (e/Å$^2$) | 60 | 60 |
| Defocus range (μM) | -1.4 – -2.6 | -1.4 – -2.6 |
| Pixel size (Å) | 1.06 | 1.06 |
| Symmetry imposed | C1 | C1 |
| Initial particles images (no.) | 1240576 | 1240576 |
| Final particles images (no.) | 93191 | 24967 |
| Map resolution (Å) | 4.2 | 5.8 |
| FSC threshold | 0.143 | 0.143 |
| **Refinement** | | |
| Map sharpening *B* factor (Å$^2$) | -143 | -156 |
| Model composition | | |
| Non-hydrogen atoms | 12459 | 12459 |
| Protein residues | 1539 | 1539 |
| Ligands | 1 (4Fe4S cluster) | 1 (4Fe4S cluster) |
| *B* factors (Å$^2$) | | |
| Protein | 149.96 | 258.16 |
| Ligand | 153.81 | 338.56 |
| R.m.s. deviations | | |
| Bond lengths (Å) | 0.008 | 0.009 |
| Bond angles (°) | 1.523 | 1.68 |
| Validation | | |
| MolProbity score | 1.96 | 2.13 |
| Clashscore | 7.69 | 12.84 |
| Poor rotamers (%) | 1.88 | 1.88 |
| Ramachandran plot | | |
| Favored (%) | 95.19 | 95.65 |
| Allowed (%) | 4.68 | 4.35 |
| Disallowed (%) | 0.13 | 0 |

**Table S2. X-ray data collection and refinement statistics**

| | **Ctf18-1-8/Pol2(1-528)** |
|---|---|
| Space group | $P3_221$ |
| **Cell dimensions** | |
| a, b, c (Å) | 126.34, 126.34, 378.43 |
| α, β, γ (°) | 90, 90, 120 |
| Resolution | 19.97 - 6.10 (6.82 – 6.10) |
| $R_{pim}$ | 0.055 (1.043) |
| $CC_{1/2}$ | 0.998 (0.362) |
| I/Iσ | 9.2 (0.8) |
| Completeness (%) | 97.2 (100) |
| Redundancy | 9.0 (10) |
| **Refinement** | |
| Resolution | 19.97 – 6.10 |
| No. reflections (free) | 8593 (840) |
| $R_{work}/R_{free}$ (%) | 27.5/33.3 |
| **No. atoms** | |
| Protein | 8613 |
| Ligand/ion | 0 |
| Water | 0 |
| **B-factors** | |
| Protein | 511.4 |
| Ligand/ion | - |
| Water | - |
| **r.m.s. deviations** | |
| Bond lengths (Å) | 0.003 |
| Angles (°) | 0.739 |
